# Circulating Biomarkers for Therapeutic Response to Immune Checkpoint Inhibitor Therapy in Patients with Advanced Lung Cancer

**DOI:** 10.1101/2025.05.21.655239

**Authors:** Kriti Jain, Deepa Mehra, Manisha Yadav, Shyam Aggarwal, Parul Chugh, Gaurav Tripathi, Surajit Ganguly, Rashmi Rana, Amit Awasthi, Nirmal Kumar Ganguly, Jaswinder Singh Maras

## Abstract

Immune checkpoint inhibitors (ICIs) have demonstrated remarkable efficacy in the treatment of advanced lung cancer (ALC), but not all patients benefit. To identify responders, patient selection, categorization, and the identification of specific predictive biomarkers are crucial. Our research aims to identify circulating metabolites that could be used to stratify patients into potential responders prior to the initiation of ICIs and to monitor clinical response in real-time during treatment. In the discovery phase, plasma samples (n = 25) from ALC patients receiving ICIs were analyzed using unbiased metabolomics and validated in a distinct patient cohort (n = 39). Five potential biomarkers were identified (FC > 1.5, p 0.05, AUC 0.81-1), including Thiourea and a combined metabolite panel (CMP) consisting of sphingosine-1-phosphate, gentisic acid, glutathione, and 4-hydroxybutanone. Early on-treatment high plasma levels of Thiourea were significantly associated with 5-year progression-free survival [Hazard ratio (HR) = 0.038, 95% confidence interval (CI) = 0.013-0.107, p<0.001] and overall survival [HR = 0.048, 95% CI = 0.019-0.120, p<0.001]. Low levels of CMP early on treatment were significantly associated with progression-free survival [HR = 8.119, 95% CI: 3.767–17.501, p<0.001] and overall survival [HR = 7.367, 95% CI: 3.517–15.433, 3.517-15.433, p<0.001]. This predictive plasma metabolite panel could serve as promising circulating biomarkers for predicting the therapeutic response of ICIs in ALC patients.

## 1. Introduction

Worldwide, lung cancer is the primary cause of cancer-related deaths, with a poor prognosis [1]. The majority of non-small cell lung cancer (NSCLC) cases are diagnosed at an advanced stage [2, 3]. NSCLC accounts for approximately 80% of all lung cancer cases.Treatment for lung cancer includes surgery, radiation therapy, chemotherapy, and more recently, ICIs [4] . In ALC, ICIs targeting programmed cell death protein 1 (PD-1) or its ligand PD-L1 have demonstrated superior efficacy to conventional chemotherapy in both the second- and first-line treatment of advanced NSCLC [5]. Currently, immune checkpoint inhibitors (ICIs) developed to treat malignant tumors, including ALC, can be classified into anti-PD-1, anti-PD-L1, and anti-CTLA-4 antibodies. Currently, ICIs developed to treat malignant tumors, including ALC, can be classified into anti-PD-1, anti-PD-L1, and anti-CTLA-4 antibodies. Amongst ICI immunotherapy agents, atezolizumab, durvalumab, ipilimumab, nivolumab and pembrolizumab are currently used as standard of care treatment for metastatic or earlier stages of cancer [6]. Presently, the Food and Drug Administration (FDA) has approved Nivolumab and Pembrolizumab, two anti-PD1 antibodies or ICIs, for the treatment of ALC [6]. However, only 25–30% of patients benefit from these treatments [7, 8]. The mechanisms underlying the variation in response patterns remain inadequately understood despite ongoing efforts. In addition, some patients suffer from severe immune-related adverse events (IRAEs), necessitating the identification of more prospective candidates for ICIs [9, 10]. To address the issues of low patient response, IRAEs, and exceptionally high costs, it is crucial to identify specific biomarkers that can stratify patients who are likely to benefit from ICIs. Currently, immunohistochemistry (IHC) staining of PD-L1 on tumour tissue specimens is the most widely used clinical biomarker for selecting patients to receive ICIs [11]. However, PD-L1 expression is not always correlated with patient survival or responses. For example, only 44.8% of PD-L1-positive patients in first-advanced-stage NSCLCs responded to ICIs, and despite PD-L1 expression, which is considered an established biomarker of response to ICIs, a substantial proportion of PD-L1-negative patients with NSCLC also experienced benefit [12, 13].

Other tissue biomarkers, such as tumour mutational burden (TMB) and microsatellite instability analysis (MSI), are also utilised in clinical practise, but they lack specificity. Blood-based immune biomarkers, neoantigens, tumor-infiltrating lymphocytes, and the diversity of the gastrointestinal microbiome have been reported to correlate with clinical outcomes [14–18]. However, these proposed biomarkers are neither precisely prognostic nor related to survival. In addition, the majority of them are based on tumour assays requiring tissue biopsies, which necessitates invasive sampling, rendering them unfeasible in the majority of cases and impractical for monitoring patient response during treatment. Recent interest has been drawn to blood-based circulating biomarkers for the prediction of response to ICIs because they are minimally invasive, requiring only blood samples, and response could also be monitored in real time [19, 20]. However, they are not yet established.

A expanding body of evidence suggests that ICIs, in conjunction with interventions targeting the metabolic pathways that obstruct anti-tumor immunity, may be a promising strategy for improving clinical efficacy and overall patient survival [19]. Recent research demonstrates that the identification of novel metabolic targets for combination therapies may be facilitated by metabolic biomarkers that guide immunotherapeutic responses. The development of mass spectrometry (MS)-based metabolomics has aided in the identification of new biomarkers for personalised treatment [21]. Notably, only a limited number of metabolomics studies have been conducted to investigate the role of circulating metabolites post-ICIs [19], the gut microbiota-derived metabolites in responsive patients [22], and the correlation between plasma metabolites and T cell-associated markers [23, 24–25]. Nonetheless, metabolic biomarkers that can accurately predict the responder (R) or non-responder (NR) status of ICIs have not yet been discovered.

In this study, plasma samples were collected to identify early metabolites that can differentiate between R and NR at baseline in ALC for significant clinical benefits. This research aims to identify a plasma biomarker panel that could stratify patients into potential R prior to the initiation of ICIs and monitor therapeutic response during treatment. Herein, on the basis of AUC values (0.8), P values (0.05), and fold change (>1.5 and 0.65), five significant metabolite biomarkers have been identified from baseline plasma samples of ALC patients. The plasma metabolite biomarker panel as a signature was observed to correlate with improved survival in patients with ALC receiving ICIs.

## 2. Materials and Methods

### 2.1 Patient selection and sample collection

ICI-treated patients with ALC (NSCLC and SCLC) older than 18 years were recruited from Sir Ganga Ram Hospital (SGRH), New Delhi. A patient cohort (n = 25) treated with ICIs as second-line therapy following chemotherapy was recruited during the discovery phase, and potential metabolite biomarkers were validated in a distinct patient cohort (n = 39) (Figure 1A). Thirty healthy volunteers were also enrolled in the investigation as a control group. All patients received ICIs: Nivolumab (200 mg, 2 l weekly) and Pembrolizumab (240 mg, 3 l weekly), and were monitored until disease progression or mortality. Magnetic resonance imaging (MRI) was used to assess the progression of the disease, and Response Evaluation Criteria in Solid Tumors Version 1.1 (RECIST 1.1) was utilized to evaluate the therapeutic response. Every 8 weeks, the clinical response to ICI treatment was evaluated. R were characterized by exemption from disease, decreased tumor size, and stable disease, while NR were characterized by an increase in tumor size or the absence of any clinical benefit within six months or less.

**Figure 1.**
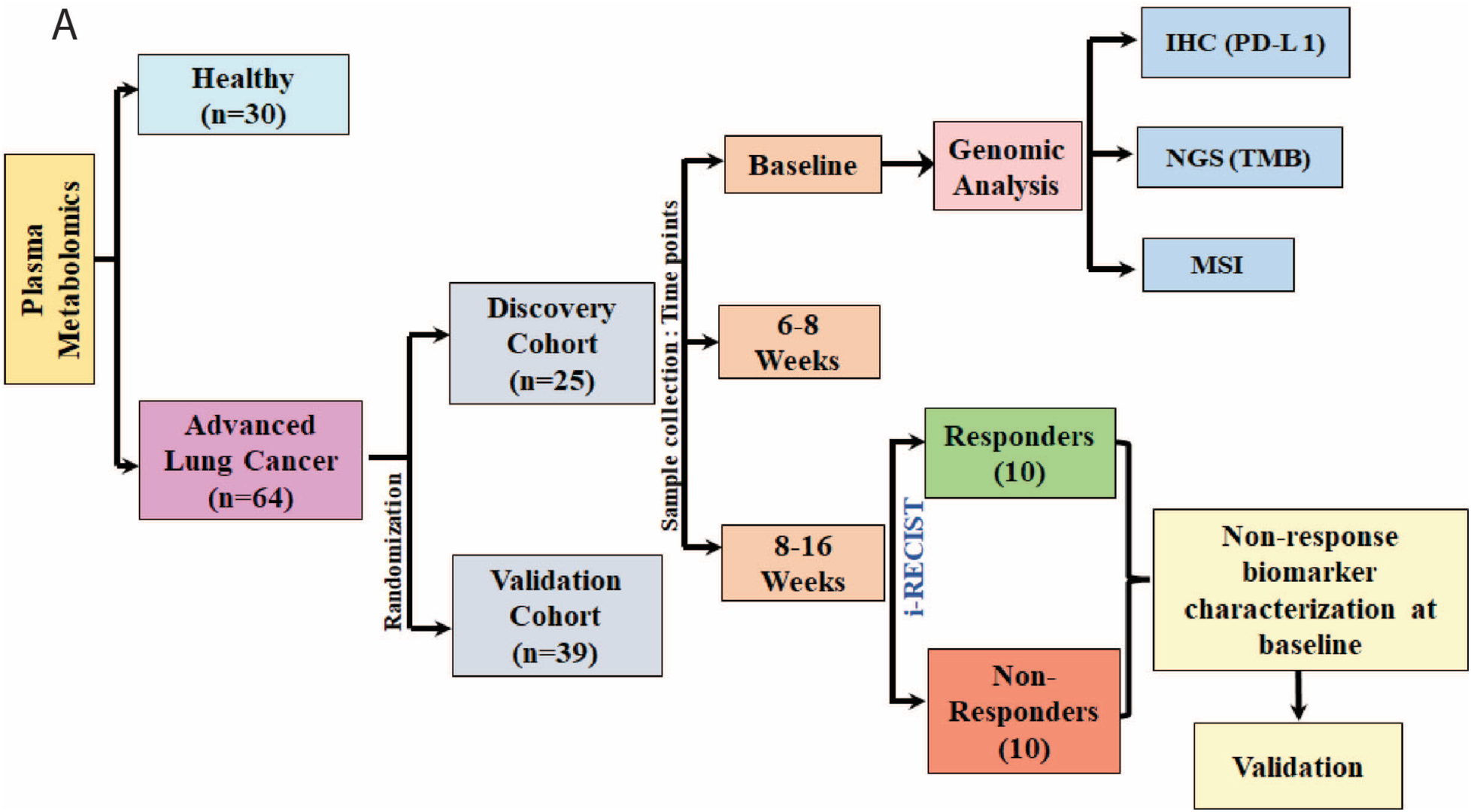
**A:** Study design. A schematic representation of the metabolomics in discovery and validation cohorts, including randomization sample collection at baseline, and time-points, along with baseline genomic analysis. Potential metabolite biomarkers are identified at baseline and validated.

Peripheral blood samples (4 ml) were collected from lung cancer patients receiving ICIs in an EDTA anticoagulation vial. Samples were collected Pre-therapy at baseline (T0) and post-treatment at two time points: 6–8 weeks (T1) and 14–16 weeks (T2). After blood was centrifuged at 3500 rpm for 15 minutes, plasma was separated and stored at -80° C until use. Formalin-fixed paraffin-embedded (FFPE) tumor tissue blocks were collected before the initiation of therapy to determine the values of established clinical biomarkers and correlate them with a response. A written consent form to participate in the study was obtained from all the patients recruited.

### 2.2 Immunohistochemistry (IHC)

Immunohistochemical staining on a formalin-fixed paraffin-embedded (FFPE) block was done to assess the expression of programmed death ligand-1 (PDL-1) in tumor tissue via the SP263 Ventana Platform. A rabbit-anti-human PD-L1/CD274 monoclonal antibody was used. Histological analysis of tumor cells (TC) and tumor-infiltrating immune cells (IC) was performed to analyze the expression of PD-L1.

### 2.3 Next-generation sequencing (NGS) for Tumor Mutation Burden (TMB)

To determine TMB, NGS (Ion Torrent, ThermoFisher Scientific) was performed using DNA extracted from FFPE tissue using the Ion Torrent Oncomine Tumor Mutation Load Assay Panel (ThermoFisher Scientific). A total of 1.65 MB were targeted, out of which 1.2 MB belong to exonic locations, covering deleterious mutations in the 43 driver genes and hotspot mutations in 61 driver genes. As per FDA and NCCN guidelines, treatment with nivolumab or pembrolizumab is beneficial for a TMB score ≥ 10.

### 2.4 Microsatellite Instability Analysis (MSI)

MSI is assessed using a panel of microsatellite markers specific to ALC, which was developed by comparing tumor tissue with normal tissue. 15 markers were multiplexed across five dye channels. PCR and fragment analysis were performed to analyze the stability and instability rate (SeqStudio Genetic analyzer, ThermoFisher Scientific). MSI-High [MSI-H] (high amount of instability in tumor >40%), MSI-Low [MSI-L] (low amount of instability in tumor, 20%), and MSS [stable] (<5% of all markers are stable) were categorized.

### 2.5 Liquid Chromatography Mass Spectrometry

An organic phase extraction protocol was used to isolate metabolites from 100 µl of the plasma sample (18). About 400 µl methanol was added to 100 µl of the plasma sample and kept overnight at -20°C for protein precipitation. These samples were centrifuged at 13,000 rpm for 10 minutes. The protein pellet was discarded, and the supernatant was isolated, dried under vacuum conditions, and reconstituted with (90:5:5) 90% water, 5% internal and external standards, and 5% acetonitrile (ACN), which was then loaded for reverse-phase chromatography in a C18 column (ThermoScientific) on an ultra-high-performance liquid chromatographic system, followed by high-resolution Q-Exactive Orbitrap mass spectrometry (MS) (ThermoScientific) (26). Identified and annotated features were subjected to log normalization and Pareto-scaling using MetaboAnalyst 5.0 and used to perform principal component analysis, partial least squares discriminate analysis, heat maps, random forest analysis, and others. Pathway enrichment patterns were analyzed using MetaboAnalyst 5.0.

### 2.6 Statistical Analysis

The results are shown as mean ± SD unless indicated otherwise. A nonparametric Wilcoxon-Mann-Whitney test Using GraphPad Prism v6 was used, and a P value < 0.05 was considered to be statistically significant. To compare variables between the two groups, the unpaired (two-tailed) Student’s *t*-test and Mann-Whitney test were performed. For comparison among more than two groups, a one-way ANOVA, Kruskal-Wallis test, and Chi-Square test were performed. Adjusted *p* values and log fold changes in LC-MS/MS data were calculated according to Benjamini and Hochberg. Expression in terms of metabolome fold changes was computed based on intensity values. Functional analysis was performed with MetaboAnalyst and KEGG pathway enrichment. Univariate statistical analysis of marker metabolites was performed with MetaboAnalyst (biomarker analysis). The biomarker model was built by binary logistic regression using a forward stepwise method. To evaluate the classification performance, receiver operating characteristic (ROC) analysis was conducted for the area under the ROC curve (AUROC). Univariate and multivariate correlations were done using SPSS version 2.6. The Kaplan-Meier method was used to analyze progression-free and overall survival. The hazard ratio (HR) was calculated from the univariate Cox Regression to determine the association between the metabolite panel and mortality. Multivariate Cox regression was done to correlate patient characteristics with the metabolite panel (SPSS v. 26).

## 3. Results

### 3.1 Patient and Randomization

Between January 2020 and March 2022, a total of 64 patients were analyzed. The patient’s clinical characteristics are listed in Table 1. The potential circulating plasma biomarkers were confirmed in two independent cohorts. Both cohorts exhibited no statistically significant differences in age, sex, nutrition, smoking status, co-morbidities, histology, disease stage, drug administered, or previous treatment. In the validation cohort, however, a significant difference was observed between R and NR in the expression of PD-L1 and MSI status.

**Table 1:** Clinical characteristics of patients in discovery and validation set.

| Characteristics | Discovery set (n = 25) |  |  | Validation set (n=39) |  |  |
| --- | --- | --- | --- | --- | --- | --- |
|  | NR(n=10) | R(n=15) | P value | NR(n=20) | R(n=19) | P value |
| **Age** |  |  |  |  |  |  |
| **Age <50** | 2 (20%) | 0 | 0.150 | 2(10%) | 0 | 0.487 |
| **>50** | 8 (80%) | 15 (100%) |  | 18(90%) | 19 (100%) |  |
| **Sex** |  |  |  |  |  |  |
| **Male** | 6(60%) | 11 (73.4%) | 0.667 | 12(60%) | 13 (68.5%) | 0.831 |
| **Female** | 4(40%) | 4 (26.7%) |  | 8(40%) | 6 (31.6%) |  |
| **Diet** |  |  |  |  |  |  |
| **Non-vegetarian** | 6 (60%) | 8 (53.4%) | 1.000 | 12(60%) | 10 (52.7%) | 0.888 |
| **Vegetarian** | 4(40%) | 7 (46.7) |  | 8 (40%) | 9 (47.4%) |  |
| **Smoking Status** |  |  |  |  |  |  |
| **Smokers** | 2 (20%) | 7(46.7%) | 0.229 | 7 (35%) | 8 (42.2%) | 0.899 |
| **Non-smokers** | 8 (80%) | 8 (53.4%) |  | 13 (65%) | 11 (57.9%) |  |
| **Histology** |  |  |  |  |  |  |
| **Non-small cell lung carcinoma(NSCLC)** | 9(90%) | 12(80%) | 0.627 | 13(65%) | 14 (73.7%) | 0.810 |
| **Small cell lung carcinoma (SCC)** | 1(10%) | 3 (20%) |  | 7(35%) | 5 (26.4%) |  |
| **Large cell carcinoma** | 1(10%) | 0 | 0.400 | 4 (20%) | 1 (5.3%) | 0.342 |
| **Squamous cell carcinoma** | 5(50%) | 7 (46.7%) | 0.870 | 10 (50%) | 8 (42.2%) | 0.863 |
| **Adenocarcinoma** | 4 (40%) | 8(53.4%) | 0.688 | 6(30%) | 10 (52.7) | 0.267 |
| **Co-morbidities** |  |  |  |  |  |  |
| **Diabetes mellitus** | 6(60%) | 10 (66.7%) | 1.000 | 9 (45%) | 11 (57.9%) | 0.628 |
| **Hypertension** | 6(60%) | 10 (66.7%) | 1.000 | 12 (60%) | 12 (63.2%) | 0.839 |
| **Pleural effusion** | 5(50%) | 9 (60%) | 0.935 | 12 (60%) | 12 (63.2%) | 0.839 |
| **Disease Stage** |  |  |  |  |  |  |
| **III stage** | 6(60%) | 10(66.7%) | 1.000 | 12 (60%) | 11 (57.9%) | 0.894 |
| **IV stage** | 4(40%) | 5 (33.4%) |  | 8 (40%) | 8 (42.2%) |  |
| **Drug administered** |  |  |  |  |  |  |
| **Nivolumab** | 8(80%) | 7 (46.7%) | 0.211 | 15 (75%) | 8 (42.2%) | 0.078 |
| **Pembrolizumab** | 2(20%) | 8 (53.4%) |  | 5 (25%) | 11 (57.9%) |  |
| **Previous Treatment** |  |  |  |  |  |  |
| **Chemotherapy** | 8(80%) | 13 (86.7%) | 1.000 | 16 (80%) | 14 (73.7%) | 0.716 |
| **Radiotherapy** | 0 | 0 |  | 1 (5%) | 3 (15.8%) | 0.342 |
| **Chemotherapy+**<br>**Radiotherapy** | 2(20%) | 2 (13.4%) |  | 3 (15%) | 2 (10.6%) | 1.000 |
| **PDL1 Expression** |  |  |  |  |  |  |
| **Highly positive (>60%)** | 0 | 5 (33.4%) | 0.061 | 0 | 14<br>(73.7%) | <0.001 |
| **Positive (30%-60%)** | 0 | 10<br>(66.7%) | 0.001 | 0 | 5<br>(26.4%) | 0.020 |
| **Negative** | 10(100%) | 0 | <0.001 | 20(100%) | 0 | <0.001 |
| **MSI Expression** |  |  |  |  |  |  |
| **MSS** | 7(70%) | 9 (60%) | 0.691 | 3 (15%) | 12<br>(63.2%) | 0.003 |
| **MSI-L** | 3(30%) | 6 (40%) |  | 17 (85%) | 7<br>(36.9%) |  |
| **MSI-H** | 0 | 0 |  | 0 | 0 |  |
| **Alcohol** | 3(30%) | 6 (40%) | 0.691 | 6 (30%) | 6<br>(31.6%) | 0.915 |
| **Family history of cancer** | 3(30%) | 5 (33.4) | 1.000 | 8 (40%) | 9<br>(47.4%) | 0.888 |
| **Weight loss** | 6 (60%) | 11<br>(73.4%) | 0.667 | 8 (40%) | 14<br>(73.7%) | 0.072 |
| **Tobbaco** | 2 (20%) | 2 (13.4%) | 1.000 | 1 (5%) | 0 | 1.000 |
| **Metastasis** | 3(30%) | 5 (33.4%) | 1.000 | 8 (40%) | 8<br>(42.2%) | 0.894 |

### 3.2 Association between expression of PD-L1, TMB, and MSI in response to ICIs

The recruited ALC patients underwent gene expression analysis. All of the patients’ levels of PD-L1, TMB, and MSI expression were evaluated. The table (online supplementary Table 1) shows the distribution of expression on the PD-L1, TMB, MSI, and tumor protein 53 (TP53) gene mutations. Figure 2A depicts an example of IHC for PD-L1 expression on FFPE tissue.

**Figure 2:**
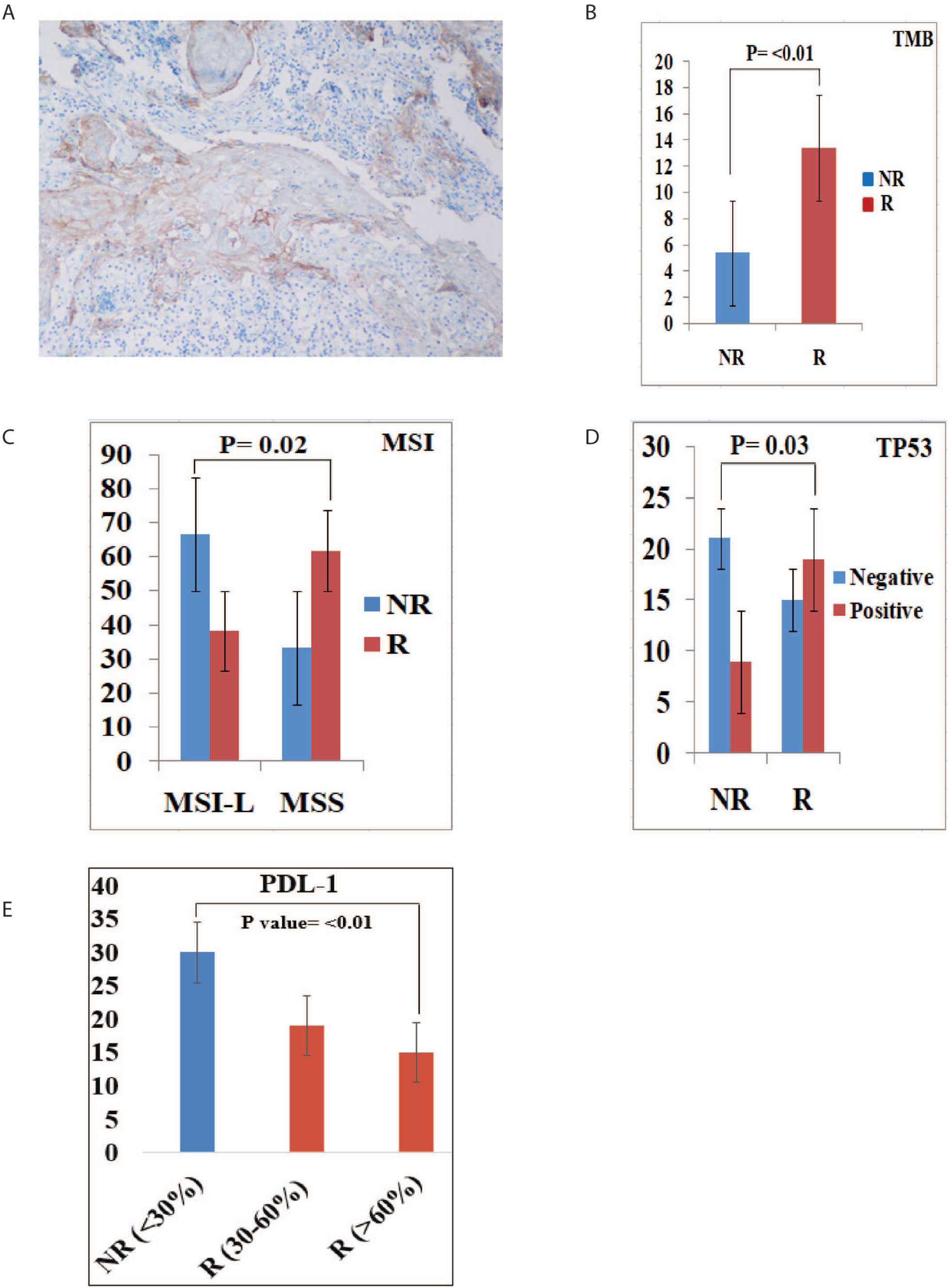
Association between expression of PD-L1, TMB, and MSI in response to ICIs. **A:** PD-L1 expression on FFPE tissue pre-ICI therapy. 20-25% of cells are positive for programmed cell death ligand 1 (PD-L1). PD-L1 staining is shown by the presence of the brown chromogen. The blue color is the hematoxylin counterstain. **B:** Mean value of Tumor Mutation Burden (TMB) between responders (R) and non-responders (NR) showing significant difference. High TMB (TMB-H) was observed in responders to immune check point inhibitor therapy. **C:** Characterization of microsatellite instability analysis (MSI) between responders (R) and non-responders (NR). **D:** Mean value of TP53 gene association between responders (R) and non-responders (NR) showing significant difference. **E:** PD-L1 expression on tumor tissue samples.

R and NR both had mean TMB values of 13.4 muts/mb (IQR 2.10-30.50) and 5.3 muts/mb (IQR 1.80-22.49), respectively. In the R subset, TMB was significantly higher (p <0.001) (Figure 2B). We found that the R subset was significantly associated with the prevalence of TP53 gene mutations (p = 0.037) (Figure 2D). No patient had microsatellite instability high (MSI-H) characterization, according to the MSI analysis; instead, R and NR were classified as MSI-low (L) and microsatellite stable (MSS), respectively. In the MSI analysis, there was a significant correlation between R and NR (Figure 2C). In R compared to NR, PD-L1 expression of more than 60% was significantly higher (p< 0.001) (Figure 2E).

### 3.3 Baseline Plasma metabolome profile of ALC patients segregates from healthy control

Plasma samples of patients receiving ICIs were subjected to an untargeted metabolomic evaluation. A total of 33441 features were identified in a positive and negative ionization mode. Out of these, 576 features were annotated and compared between the study groups (Online Supplementary Table 2A). These metabolites broadly belong to classes of amino acids, fatty acids, and lipids. Plasma samples of healthy control and ALC patients were subjected to untargeted metabolomics evaluation. A total of 473 differentially expressed metabolites (DEMs; FC > 1.5, p 0.05) (Online Supplementary Table 2B), was identified from the baseline analysis of the metabolome profiles of ALC and healthy control. Out of these, 14 were downregulated, and 459 had substantial upregulation (Figure 3A). The principal component analysis (PCA) a (Figure 3B), along with hierarchical clustering analysis, successfully distinguished baseline lung cancer metabolomes from healthy controls (Figure 3C), as well as a summary of the most important metabolite (VIP; varied importance in projection) characteristics as shown in (online supplementary Figure A).

**Figure 3:**
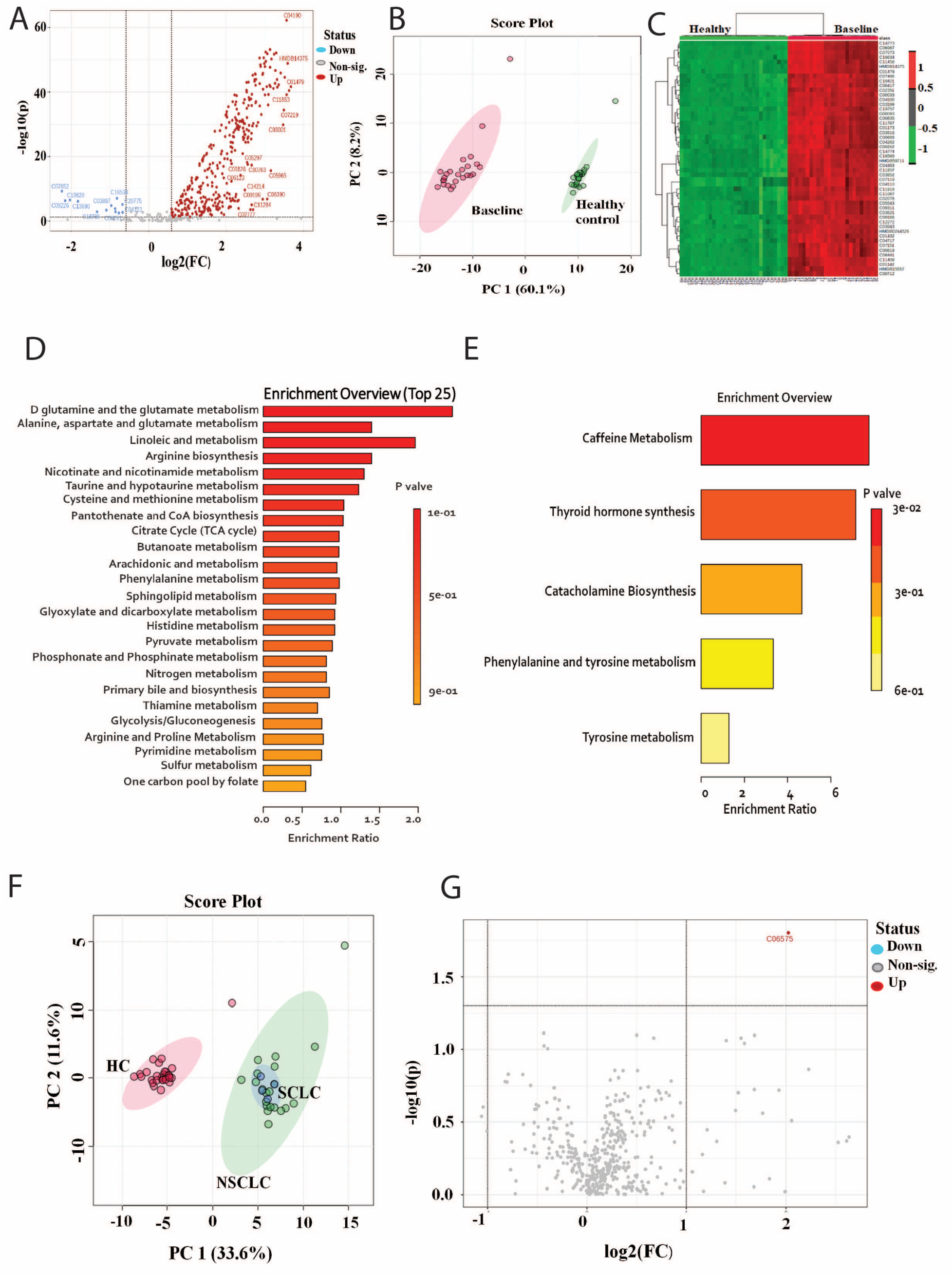
Baseline Plasma metabolome profile of ALC patients segregates from healthy control. **A:** Volcano plot showing differentially expressed metabolites of lung cancer patients at baseline as compared to healthy control, along with their expression status (Red dots = upregulated, blue=downregulated and grey = non-significant metabolites) **B:** Principal component analysis (PCA) showing clear segregation profiles of lung cancer patients at baseline as compared to healthy control, whereas the Pink dots correspond to baseline lung cancer and green dots correspond to healthy control. **C:** Heat map and hierarchical clustering analysis of top 50 metabolites shows differential expression of metabolites between baseline lung cancer pateints (red bar) from healthy control (green bar). The expression is given as red = upregulated, green = downregulated, and black = unregulated. **D:** KEGG pathway analysis of upregulated metabolites in baseline lung cancer as compared to healthy control. **E:** KEGG pathway analysis of downregulated metabolites in baseline lung cancer as compared to healthy control. **F:** The principal component analysis (PCA), shows that the metabolic profile of NSCLC overlapped with the metabolic profile of SCLC when compared to healthy control. **G:** Volcano plot showing differentially expressed metabolites at NSCLC as compared to SCLC, along with their expression status (Red dots = upregulated, blue=downregulated and grey = non-significant metabolites).

Metabolites associated with D-Glutamine, D-glutamate metabolism, Alanine, Aspartate metabolism, and others were found to be increased in ALC patients (Figure 3D). Similarly, downregulated metabolites were associated with caffeine metabolism and thyroid hormone synthesis, and others (Figure 3E).

Additionally, a subgroup analysis was conducted to compare individuals with non-small cell lung cancer (NSCLC), small cell lung cancer (SCLC), and a control group consisting of healthy individuals. The results of the principal component analysis (PCA) indicate that the metabolic profile of non-small cell lung cancer (NSCLC) exhibits similarities with the metabolic profile of small cell lung cancer (SCLC) in comparison to a healthy control group (Figure 3F). However, a subsequent comparison was conducted between the NSCLC and SCLC groups. The findings from the Partial Least Squares Discriminant Analysis (PLSDA) indicate that the metabolic profile exhibited distinct characteristics (Supplementary Figure B). However, there was no statistically significant difference observed in the expression of metabolites between the two groups (Figure 3G).

### 3.4 The baseline plasma metabolome profile robustly differentiates R and NR

A total of 214 differentially expressed metabolites (DEMs) were identified (FC > 1.5, *p* < 0.05) (Figure 4A). Amongst these, 188 were significantly upregulated, and 26 were downregulated in NR compared to R. Based on the results of PCA (Figure 4B) and a heat map showing hierarchical clustering analysis, NR and R were considerably discriminated (Figures 4C), and variable importance in the project (VIP) traits were determined (Figure 4D). ALC plasma metabolites that were increased in NR were associated with sphingolipid metabolism as well as D-glutamine and D-glutamate metabolism (Figure 4E). A comparison of the baseline metabolome profiles of NR and R revealed a distinct increase in sphingolipid metabolism, taurine and hypotaurine metabolism, and D-glutamine and D-glutamate metabolism. Thus, an increase in sphingolipid and amino acid metabolism has extraordinarily broad effects on lung cancer cells and has also been reported to accelerate and promote proliferation and non-dependent growth, thereby increasing tumor progression. Downregulated metabolites were related to arginine, proline metabolism, and the urea cycle in R (Figure 4F).

**Figure 4:**
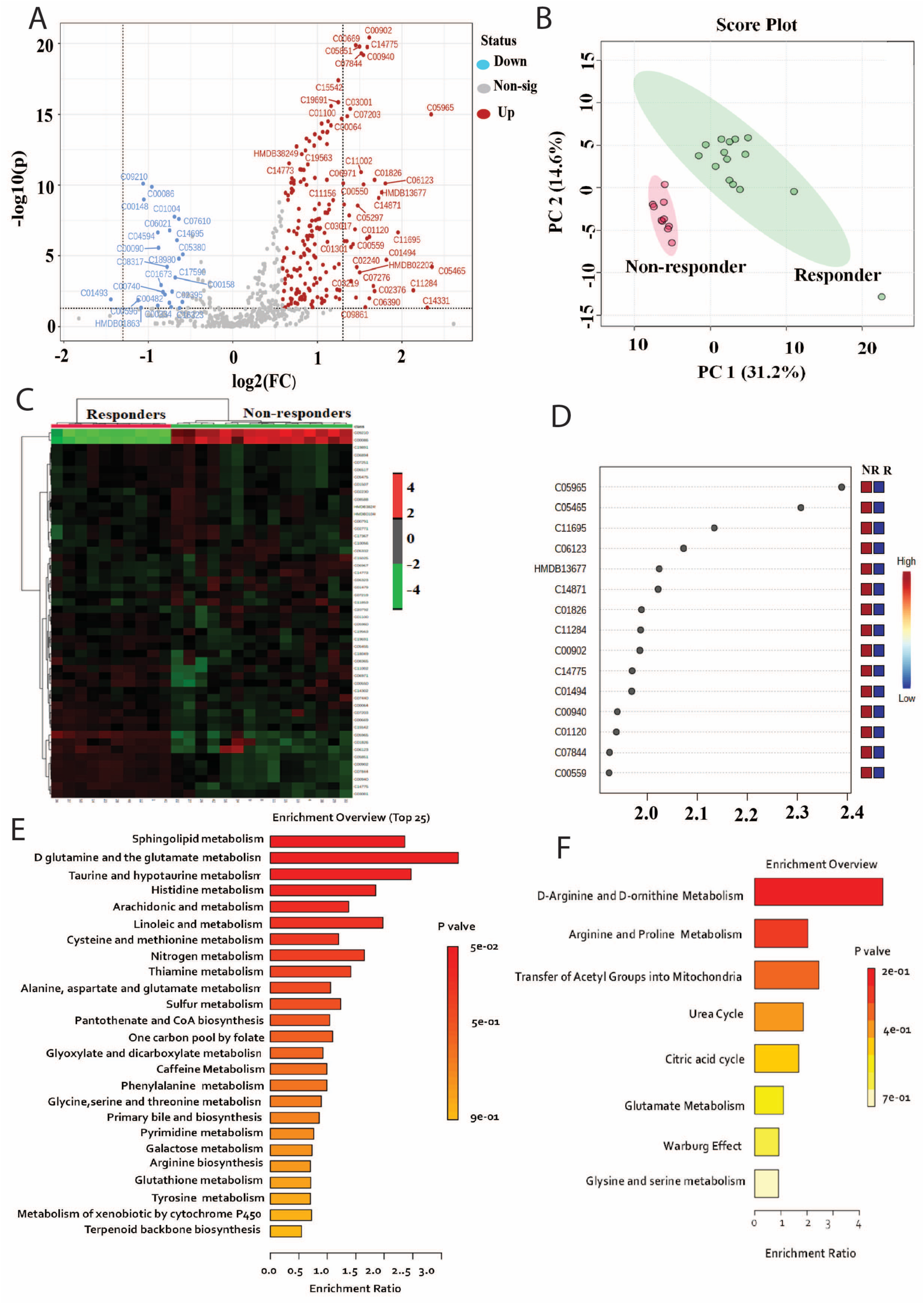
The baseline plasma metabolome profile robustly differentiates R and NR. **A:** Volcano plot showing differentially expressed metabolites of lung cancer patients in Non-responders as compared responders at baseline, along with their expression status (Red dots = upregulated, blue=downregulated and grey = Non-significant metabolites.) **B:** Principal component analysis (PCA) showing clear segregation of metabolome profiles of lung cancer patients in Non-responders as compared responders at baseline, where the Pink dots correspond to non-responders, and green dots correspond to responders. **C:** Heat map and hierarchical clustering analysis of top 50 metabolites shows differentially expression of metabolites between non-responder lung cancer patients (red bar) from responders (green bar). The expression is given as red = upregulated, green = downregulated, and black = unregulated. **D:** Variable importance in projection (VIP) Plot showing top 15 most important metabolite features identified in non-responder from responder at baseline lung cancer patients. **E:** KEGG pathway analysis of upregulated metabolites in non-responders and responders at baseline lung cancer. **F:** KEGG pathway analysis of downregulated metabolites in non-responders and responders at baseline lung cancer.

### 3.5 Weighted metabolome correlation network analysis (WMCNA) corresponding to the response to ICIs in ALC patients

To gain insights into the temporal changes that occurred in the metabolic profile upon administration of ICIs in R and NR, WMCNA was performed. WMCNA was used on the metabolic profiles of R and NR at T0, T1, and T2 to find metabolic modules that are unique to R and NR. These are groups of metabolites whose expression changes in the same way over time. WMCNA clustered 576 metabolites into 12 modules (Figure 5A). The heat map shows how, after therapy, many metabolites in R were turned on or off, while in NR, the expression of metabolites didn’t change as much as it did in R. Figure 5B shows how a module-trait relationship was used to find the R and NR-specific modules at baseline (the phenotypes of NR and R at T0, T1, and T2 are called traits). Module trait relationships identified the ’Green Yellow’ module and the Blue’ module specific to R (up-regulated at baseline) (Figure 5C). The plot represents the expression of metabolites in the ’Green Yellow’, and ‘Blue’ modules, which are linked to Phenylalanine, tyrosine metabolism, tryptophan biosynthesis, etc., and Sphingolipid metabolism, Pantothenate biosynthesis, etc., respectively. Thus, boosting sphingolipid and amino acid metabolism has extremely broad impacts on lung cancer cells and has also been observed to accelerate and enhance proliferation and non-dependent growth, thereby promoting tumor development. The purple module specific to NR (up-regulated at baseline) is significantly linked to caffeine, nicotinate, and nicotinamide metabolism, Glycerolipid metabolism, and the citrate cycle (TCA cycle) (Figure 5D) (Figure 5E). Various studies have examined the relationship between coffee consumption and the risk of developing cancer. Coffee use in particular has reportedly been linked to an increased risk of lung cancer [28]. The metabolism of amino acids helps cancers survive and also drives their growth. Since glutamine is a key source of nitrogen and carbon for the synthesis of amino acids and for the renewal of the TCA cycle, it is essential for the survival of many tumor cells [27]. Hence, WMCNA analysis showed that studying these clusters of metabolites and their associated pathway analysis temporally may help monitor responses to ICIs.

**Figure 5:**
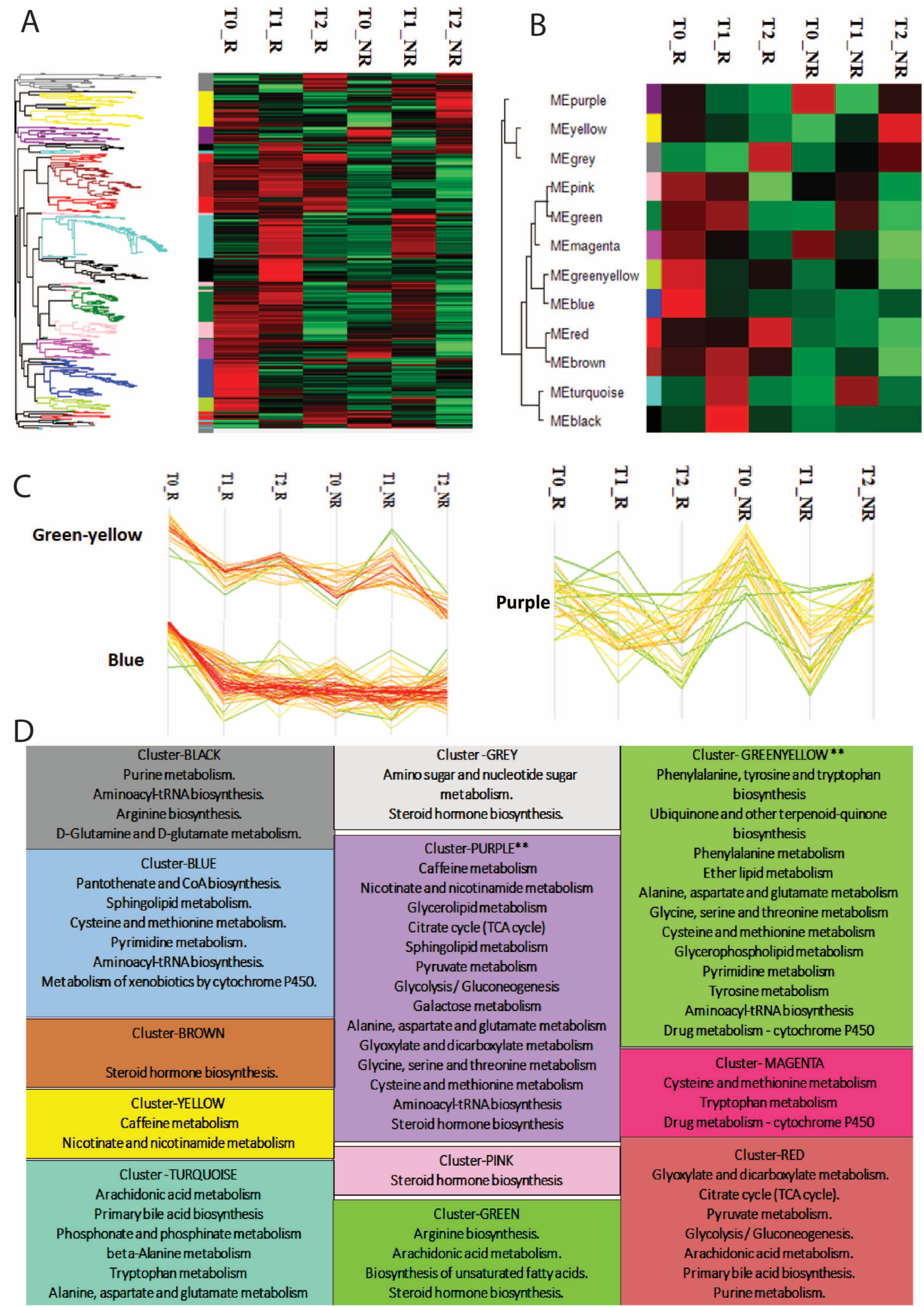
Weighted metabolome correlation network analysis (WMCNA) corresponding to the response to ICIs in ALC patients. **A:** Weighted metabolome correlation network analysis (WMCNA) Heat map showing clustered 576 metabolites into 12 modules on the basis of similar trend in their expression over time specific to R and NR at different T0,T1,T2 time points. **B:** Module trait relationship Heat map showing the correlation between modules and traits (T0, T1, T2 are referred to as a trait) specific to R and NR. The more intense the box color, the more negatively (green) or positively (red) correlated is the module with the trait. **C:** Module trait relationship graph-plot represents the expression of metabolites at different time points in the ’Green Yellow’ module and ‘Blue’ module specific to R (up-regulated at baseline). **D:** Module trait relationship graph-plot represents the expression of metabolites at different time points in ’Purple’ module specific to NR (up-regulated at baseline). **E:** Panel showing most altered pathways from each color modules from which Green yellow and purple are significantly associated pathways.

### 3.6 Identification of plasma metabolite biomarker panel predicting response to ICIs at baseline

All DEMs at baseline between R and NR were determined using a volcano plot, and 35 significant metabolites were identified using fold change (>1.5), p-value (<0.05), and AUC value ranging between 0.87-1.09. Out of these 35 significant metabolites, we identified five potential biomarker panels by confirming them with authentic standards and online literature (Figure 6A). Based on these, the five highly significant plasma metabolites obtained were distinguishing R and NR. These were – Thiourea (AUC -0.99, FC-0.46), sphingosine 1-phosphate (AUC-1, FC-4.43), gentisic acid (AUC-0.93, FC-3.03), glutathione (AUC-0.97, FC-2.23) and 4-hydroxy butanone (AUC-0.99, FC-2.53) and the P value of all biomarker were (p<0.001). R showed significantly higher levels of thiourea whereas the other four significant metabolites namely Sphingosine 1-phosphate, gentisic acid, glutathione, and 4-hydroxy butanone were significantly downregulated, termed here as combined metabolite panel (CMP) (Figure 6B, Figure 6C).

**Figure 6:**
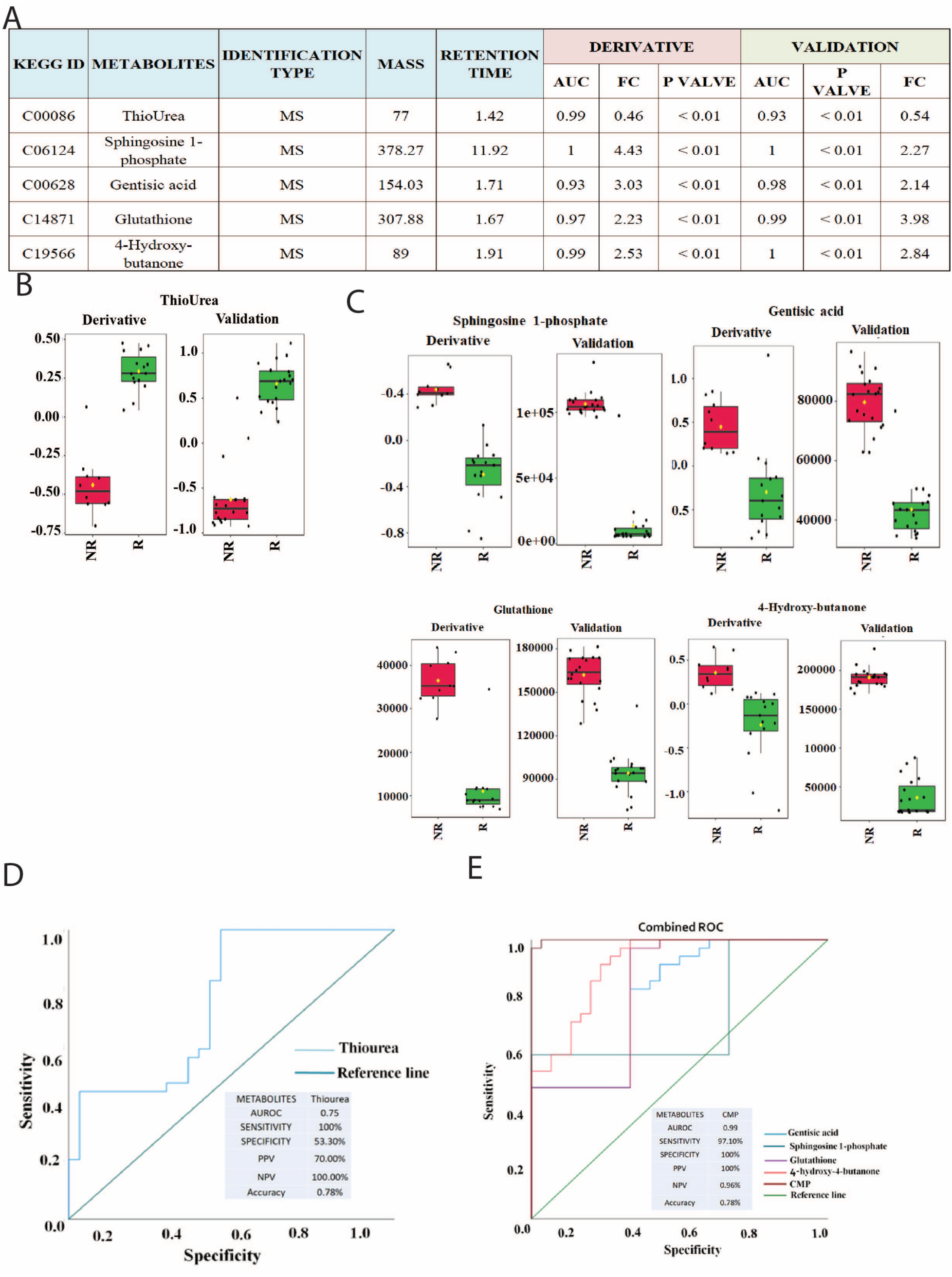
Identification of plasma metabolite biomarker panel predicting response to ICIs at baseline. **A:** Plasma metabolite biomarker panel showing the identification type for each biomarker associated from discovery and validation set. **B:** Plasma levels of Thiourea at early on-treatment in responders and non-responders of the discovery and validation sets. The box plots depict the minimum and maximum values (whiskers), the upper and lower quartiles, and the median. **C:** Plasma levels of potential marker metabolites Sphingosine 1-phosphate, Gentisic acid, Glutathione ,4-Hydroxy-butanone at early on-treatment in responders and non-responders of the discovery validation sets. The box plots depict the minimum and maximum values (whiskers), the upper and lower quartiles, and the median. **D:** Receiver operating characteristic (ROC) analysis Thiourea in the discovery and validation set. **E:** Receiver operating characteristic (ROC) analysis of CMP in the discovery and validation set.

The receiver operating characteristic (ROC) analysis shows Thiourea and CMP distinguishing R and NR (Figure 6D, Figure 6E). The cutoff value for the R for Thiourea and CMP used was (log>=10.57) and (log<=14.44) respectively. Based on this the area under the curve (AUROC) for Thiourea was 0.755 [95% confidence interval (CI), 0.633-0.876], with a sensitivity of 100% and specificity of 53.3%, and for CMP, AUROC was 0.999 [95% CI, 0.996-1.000] with a sensitivity of 97.1% and specificity of 100%.

### 3.7 Correlation between plasma metabolite biomarker panel and clinical-pathologic characteristics

The biomarker panel was correlated with the relevant clinical data remarkably capable of distinguishing R and NR (Figure 7) (R^2^<0.5). Thiourea metabolite showed a significant positive correlation with PDL1 expression in R whereas CMP was downregulated in R which correlated with no PDL1 expression. PDL1 expression is an FDA-approved biomarker for lung cancer patients being treated with ICIs [29]. We observed that our clinical plasma metabolite panel positively correlated with PDL1 expression.

**Figure 7:**
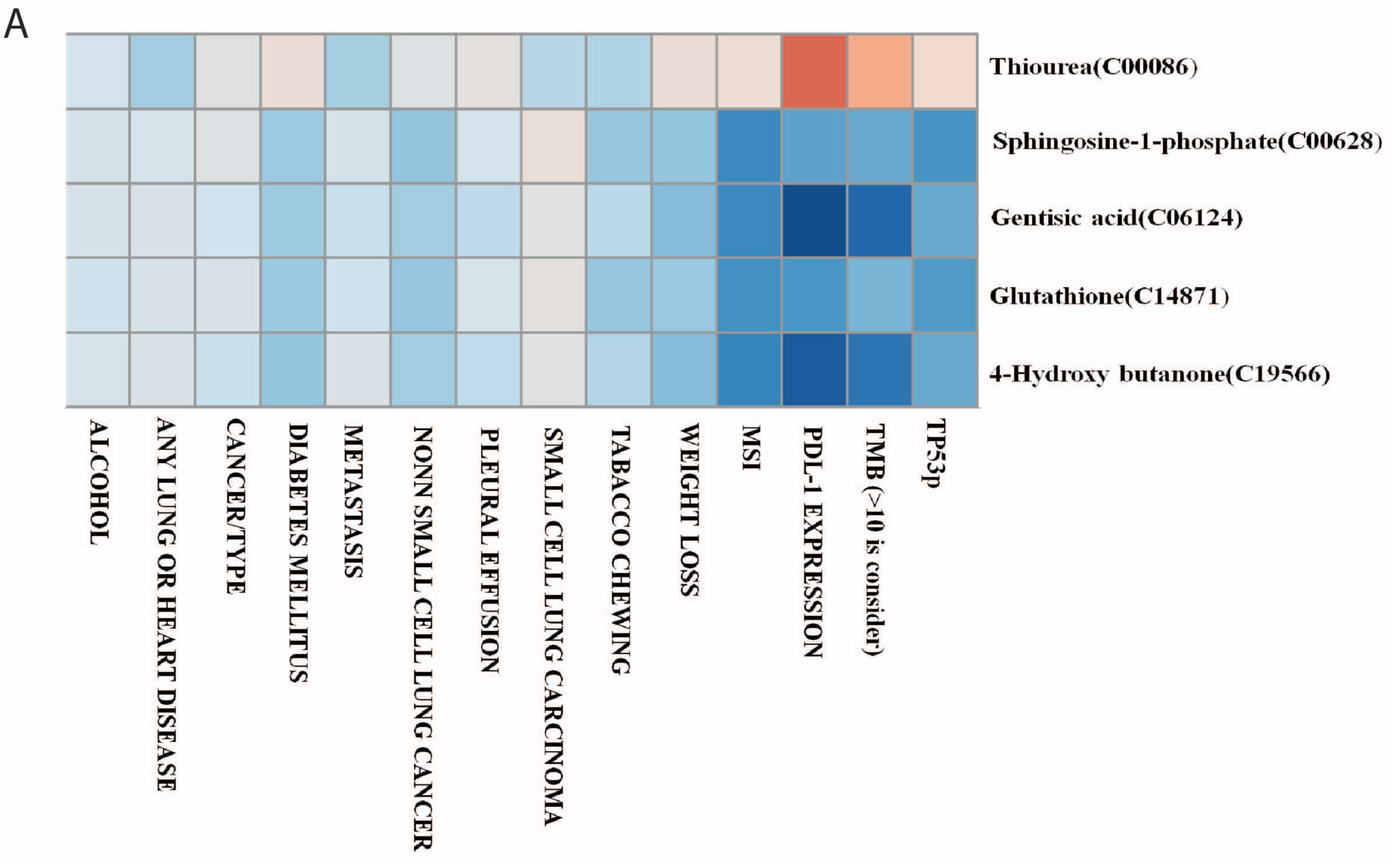
Plot showing correlation between Clinical parameter of cancer patients with associated biomarker panel.

### 3.8 Plasma metabolite biomarker panel associated with improved patient survival in discovery and validation

The potential metabolite biomarkers were evaluated in two independent cohorts. We determined the absolute concentration of thiourea and CMP in plasma, and the measurements were reproducible in the validation set also. The logistic regression model of the metabolite panel associated with response to ICIs was constructed. The association between thiourea and CMP with the clinical outcome of the patients receiving ICIs was observed. The plasma levels of variables thiourea and CMP at T0 were significantly associated with progression-free survival (PFS) [HR = 0.038, 95% CI, 0.013-0.107, p<0.001; HR=8.119, 95% CI, 3.767-17.501, p<0.001, respectively] (Figure 8A).

**Figure 8.**
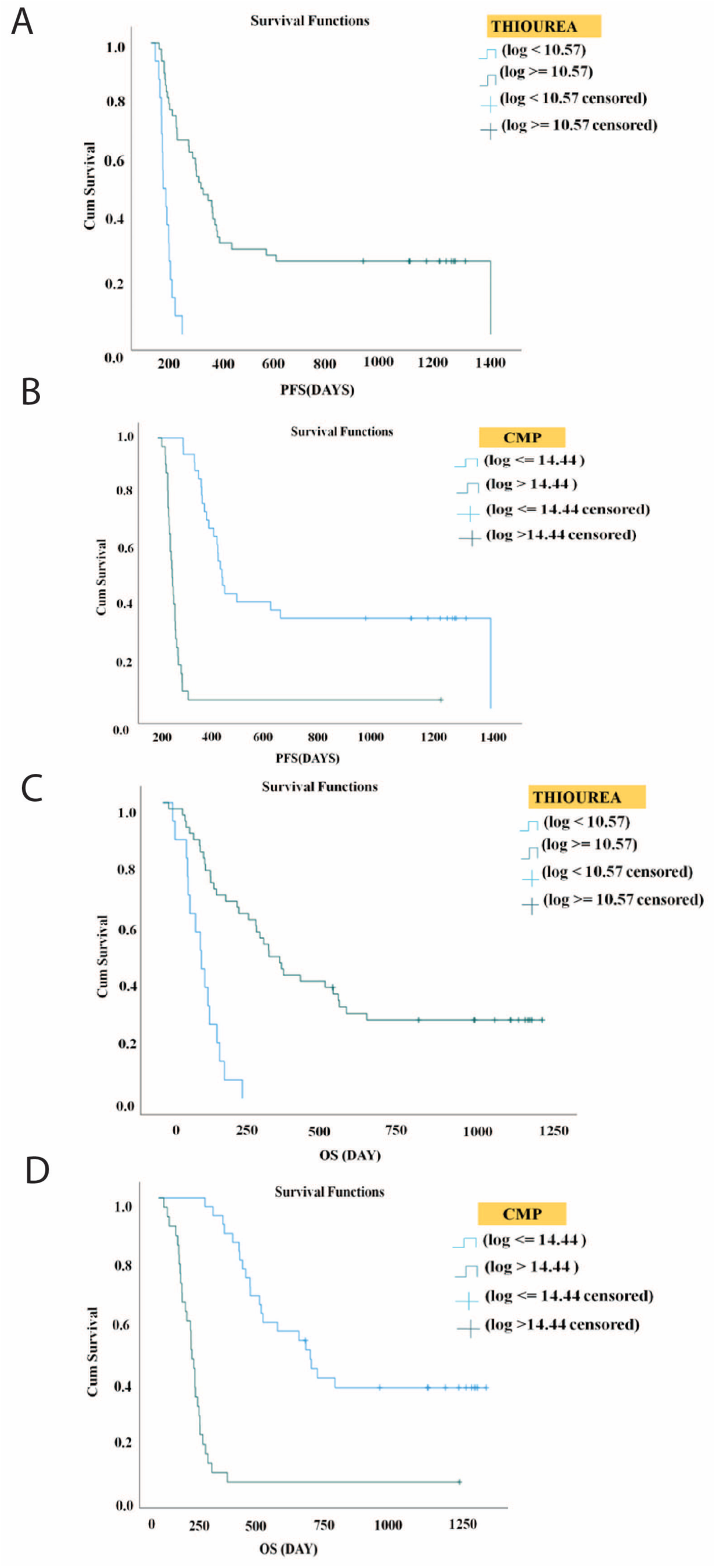
**A:** Kaplan-Meier analysis for progression-free survival in lung cancer patients by plasma level of Thiourea metabolites in discovery and validation set. **B:** Kaplan-Meier analysis for progression-free survival in lung cancer patients by plasma levels of CMP metabolites in discovery and validation set. **C:** Kaplan-Meier estimates of overall survival in lung cancer patient by plasma level of Thiourea in the discovery and validation set. **D:** Kaplan-Meier estimates of overall survival by plasma levels of CMP in the discovery and validation set.

The median PFS of patients with high levels of thiourea (log>=10.57) was 205 days (95% CI, 130-280 days, p<0.001) and low levels of CMP (log<=14.44) were 270 days (95% CI, 236-304 days, p<0.001) respectively (Figure 8B). These were the significant predictors of PFS in the univariate analysis and did not remain as the independent predictors in the multivariate analysis.

The levels of thiourea strongly correlated with overall survival (OS) (HR= 0.702, 95% CI, 0.328-1.501, p=0.361) whereas CMP was significantly associated with OS (HR=7.833, 95% CI, 3.806-16.123, p<0.001). In the univariate analysis, patients with high levels of thiourea (log>=10.57) had longer OS (median OS of 362 days, 95% CI, 265-459 days) than patients with low levels of thiourea (log<10.57) (median OS of 128 days, 95% CI, 89-167 days) (Figure 8C).

Also, patients with low levels of CMP (log <=14.44) have longer OS (median OS of 583 days, 95% CI, 374-792 days) as compared to patients with high levels of CMP (log >14.44) (median OS of 128 days, 95%, 98-157 days) (Figure 8D). In the multivariate analysis, PD-L1 positive expression (>60%) remained the independent predictor of OS (HR=0.018, 95% CI, 0.005-0.063, p<0.001). Hence, thiourea and CMP at T0 were identified as the potential biomarkers predictive of clinical outcomes in patients with ALC.

## 4. Discussion

In an era of ALC and other cancer therapies, ICIs targeting PD-1, PD-L1, and cytotoxic T-lymphocyte antigen 4 (CTLA-4) have achieved tremendous success; however, only a subset of patients can obtain benefits [30]. Development of robust predictive biomarkers that can both predict and monitor therapy response remains the current challenge. Currently, in clinical practice, patient selection based on the expression of PD-L1 by IHC and TMB is used to improve the prediction of response to ICIs in ALC. In KEYNOTE-001 and KEYNOTE-024, PD-L1 tumor proportion score (TPS) ≥ 50%, treatment with pembrolizumab treatment lengthened the survival time of NSCLC patients, making it a suitable marker for guiding the selection of patients for ICIs [31]. However, other studies have shown that patients have responded well to therapy regardless of PD-L1 expression, highlighting the limitations of PD-L1 as an independent predictor of response [32].Tumor heterogeneity and complex dynamism of immune escape mechanisms are yet not fully understood and emphasize the need to develop a set of specific biomarkers. However, in the past decade, circulating biomarkers have emerged as significant surrogates for disease monitoring in cancer.

Based on metabolomic profiling, we have identified a plasma metabolite panel that can aid in predicting response to ICIs in patients with ALC. Thiourea and CMP were identified and validated as predictive biomarkers for the clinical outcomes of ICIs in an independent patient cohort. This is the first report to our knowledge of validated plasma metabolite biomarkers predictive of response to ICIs in ALC. In addition, this blood-based test for these biomarkers is less invasive than tissue biopsies, which are also unavailable in some patients, and samples can be collected at multiple time points for monitoring the patient’s progress.

The plasma biomarker panel identified in this study not only has the potential to predict response to therapy, but also provides novel insights into the mechanism of resistance, suggesting metabolic interventions/drugs that could be administered in conjunction with ICIs to maximise therapy response outcome and improve clinical efficacy.

Thiourea compound is often used in the treatment of cancer.[new 33 and 34]. Previous research has identified multiple derivatives of 1-benzoyl-3-methyl thiourea that exhibit anticancer properties [new 35]. In previous investigations, it has been observed that the utilization of thiourea derivatives in conjunction with complexed platinum can lead to an augmentation of anticancer efficacy. The enhancement of the anticancer efficacy of thiourea can be achieved through structural alterations, such as the creation of a platinum (II)-thiourea complex [new 36]. A study also assessed thiourea complexes as anticancer drug candidate for lung cancer and concluded it to be a suitable anticancer agent [new 37]. Based on the aforementioned information, further studies may employ molecular docking and molecular dynamics techniques to facilitate the development of drug candidates in silico, targeting thiourea compound for ALC.

Thiourea and urea are regarded as privileged structures within the field of medicinal chemistry. These moieties collectively form a fundamental structure found in numerous drugs and bioactive compounds, which possess diverse therapeutic and pharmacological properties. The urea cycle (UC) converts excess nitrogen into urea, a nitrogenous compound that can be discarded [old 33]. UC dysregulation is a prevalent metabolic phenomenon in cancer that generates mutational bias (purine-to-pyrimidine ratio) and neopeptides, thereby worsening the patient’s prognosis and enhancing the response to checkpoint inhibitor therapy independent of PD-L1, TMB, and MSI. In a study, R of ICIs exhibited significantly higher UCD and mutational bias than NR. Silencing of the UC enzyme ASS1 promotes cancer proliferation by diverting the substrate aspartate to the CAD enzyme, thereby generating neopeptides and favouring pyrimidine synthesis over urea disposal [old 34]. As a therapeutic intervention, silencing of ASS1 with the goal of increasing UCD levels to improve the efficacy and coverage of ICIs can be targeted.

In our findings, we found that baseline plasma Thiourea levels were significantly higher in the R group compared to the NR group, which suggests a significant role for Thiourea in exhibiting anticancer propertie. Patients with high levels of Thiourea may be selected as a potential candidate for ICIs.

Sphingosine-1-phosphate (S1P) promotes tumor cell proliferation, chemotherapy, radiotherapy, and immunotherapy resistance, and metastasis. Previous studies have demonstrated S1P’s role in the regulation of cancer cell death and drug resistance. Dysregulation of sphingolipids such as sphingosine-1-phosphate (S1P) has been linked to drug resistance in breast, melanoma, and colon cancers [35, 40]. Multiple genome-wide studies have identified sphingolipid metabolism as one of the key dysregulated pathways in lung cancer patients, strongly indicating that harnessing and restoring sphingolipid homeostasis in patients may be an effective therapeutic strategy for treating the disease [36]. The expression of B3GNT5/GAL3ST1 (sphingolipid metabolic enzymes) controls the lacto/neolacto-series glycosphingolipid/sulfatide balance as a checkpoint for sphingolipid metabolic reprogramming during tumour progression, according to a study by Meng et al [37]. In a separate study by Imbert et al., sphingosine kinase-1 (SK1) was identified as a key regulator of anti-tumor immunity, and increased expression of SK1 in tumour cells was found to be significantly associated with shorter survival in metastatic melanoma patients treated with immunochemotherapeutic agents (ICIs). They demonstrated that SK1 downregulation increases the efficacy of ICI. Targeting SK1 increases responses to ICIs in murine models of melanoma, breast, and colon cancer; therefore, silencing SK1 decreases the expression of various immunosuppressive factors in the tumour microenvironment to limit T regulatory cell (Treg) infiltration [38]. Alternatively, high SK1 expression in melanoma cells was associated with diminished ICI efficacy [39]. Lung adenocarcinoma cells were observed in some studies to have increased EGFR expression and proliferation, thereby promoting tumour progression. Our findings indicate a significantly higher level of sphingosine-1-phosphate in NR compared to ICIs, suggesting that a change in plasma sphingolipidomes may improve the efficacy of ICIs. Consequently, the management of sphingolipids and S1P metabolism may be deemed important for enhancing the efficacy of ICIs. Some active clinical trials that target sphingolipid metabolism to activate pro-cell death ceramide signalling and/or inhibit pro-survival are undergoing evaluation. Current research on S1P signalling utilising immunological, genetic, or pharmacological tools provides novel strategies for the development of immunochemotherapeutic agents (ICIs) for various types of cancer. Together, these mechanisms by which sphingolipids regulate the signalling of cancer cells will aid in the improvement of future immunotherapy strategies [40]. 4-hydroxy butanone is a proposed biomarker for tobacco smoke exposure, which causes cellular stress, and the releasing adducts are formed by metabolic activation of N’-nitrosonornicotine and 4-(methylnitrosamino)-l-(3-pyridyl)-l-butanone [41]. Its function in cancer is unknown. Gentisic acid (GA) is a minor aspirin catabolite that endogenously generates a siderophore for the transport of iron, which stimulates tumour growth. Several studies have indicated that GA has the potential to be an anti-cancer adjuvant capable of inhibiting the formation of glioma cell colonies [42]. However, its role in ICIs has never been investigated. In accordance with our findings, GA levels were significantly elevated in NR, which may suggest malfunctioning iron transport that led to the proliferation of tumour cells. As a potential biomarker candidate, GA deserves to be studied in ALC patients receiving ICI in the future.

Our results also indicate that Glutathione (GSH) metabolite levels are significantly elevated in NR, and its metabolism is known to be involved in cancer progression and treatment resistance. Previous research has shown that excess GSH promotes tumour progression, detoxifies xenobiotics, maintains adequate cysteine levels, and confers therapeutic resistance on cancer cells [43]. This indicates that targeting GSH utilization/synthesis with prodrugs such as Quinone methide (QM) prior to the administration of ICIs can improve their therapeutic efficacy [43,44] and [44,45]. In a recent study [45,46], glutathione peroxidase was found to be significantly related to the efficacy of ICI. We also found Glutathione metabolism to be significantly elevated in NR, which is associated with arachidonic acid metabolism and neutralisation of reactive oxygen species, which likely decreases intracellular inflammation by inhibiting NF-B signalling and extracellular activation of host immunity, thereby reducing ICI response despite high PD-L1 expression and TMB. In contrast to established clinical tissue-based biomarkers, such as PD-L1, TMB, and MSI, Thiourea and CMP appear to play a significant role in predicting response.

The present investigation had some limitations. The metabolite exhibits a high degree of dynamism and susceptibility to many circumstances. Therefore, it is imperative to validate further the findings of this study in order to ensure their consistency and reproducibility. Furthermore, it is worth noting that a subset of patients in the study were deemed ineligible for surgical management prior to undergoing ICIs. This particular aspect introduces the possibility of selection bias, which could potentially impact the outcomes observed. In addition, it is recommended that future research endeavours focus on conducting bigger sample sizes, multi-center collaborations, and prospective studies in order to validate the metabolomics nomogram. This will ensure the provision of robust and dependable data for the potential therapeutic application of this approach.

In conclusion, our study demonstrated that elevated Thiourea levels in conjunction with a CMP composed of Sphingosine-1-phosphate, gentisic acid, glutathione, and 4-hydroxy butanone may serve as a potential biomarker panel predictive of clinical response to ICIs in ALC. Targeting these metabolites could be an attractive strategy for improving response and aiding in patient stratification. The discriminant metabolomics profiling developed in this study presents a viable and easy approach for tailoring treatment to individual patients. The exceptional precision of metabolomics profiling has the potential to forecast the impact of chemo-immunotherapy in patients with ALC. However, the results must be validated in a larger patient cohort to serve as an effective therapeutic decision tool.

## Data Availability Statement

The original contributions presented in the study are included in the article/ Supplementary Material. Further inquiries can be directed to the corresponding author.

## Ethics Statement

The study involving human participant was reviewed and approved by “Sir Ganga Ram Hospital Ethics Committee” (No. EC/04/19/1499; Approval date: 02/07/2019). All the patients provided their written informed consent form and willingness to participate in this study and for the publication of any potentially identifiable images or data included in this article.

## Author Contributions

All authors listed have made a substantial, direct, and intellectual contribution to the work and approved it for publication.

## Acknowledgement

We would like to acknowledge the patients and their families for cooperation in providing us the samples being used in this study. We are also thankful to Dr. Lal Path Labs National Research Laboratory for providing us the technical support in performing the genomics experiments being conducted in this study.

## Supplementary Material

The supplementary material for this article can be found online.

