## Supplementary Figure 1 for "Circulating Biomarkers for Therapeutic Response to Immune Checkpoint Inhibitor Therapy in Patients with Advanced Lung Cancer"

**SUPPLEMENTERY FIGURES**

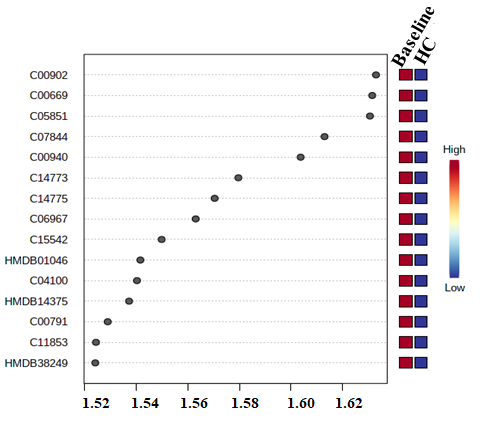
**Supplementary Figure 1:** Variable importance in projection (VIP) Plot showing top 15 most important metabolite features identified at baseline lung cancer pateints vs healthy control.

**Supplementary Figure 2:** The Partial least square discriminant analysis (PLSDA) plot showing the clear distinguished, metabolomic profiles of NSCLC and SCLC.

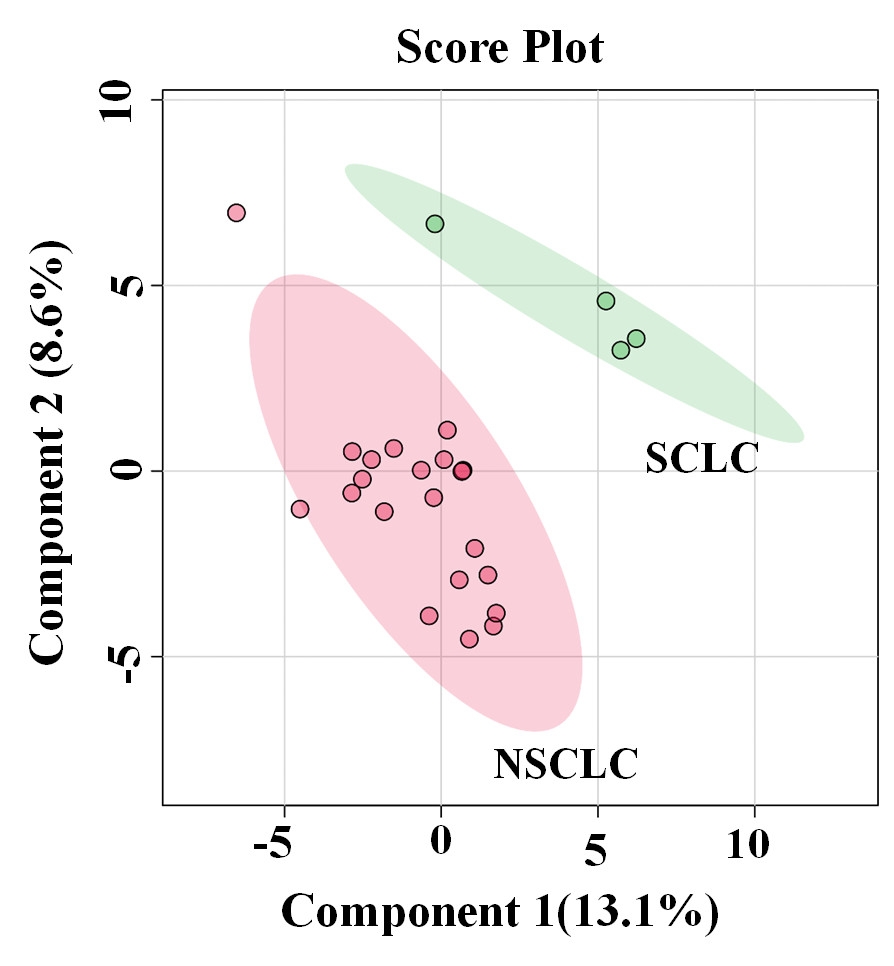

**Supplementary Figure 3:**

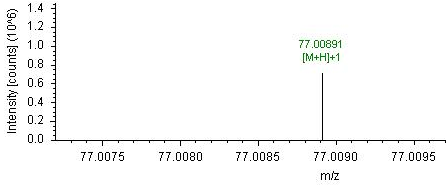
**Figure 3A:** Fragmentation pattern of thiourea leading to signal positive intensities ions [M+H]+1 present in the mass spectrum.

**Figure 3B:** Fragmentation pattern of Sphingosine 1-phosphate leading to signal negative intensities ions [M-H]-1 present in the mass spectrum.

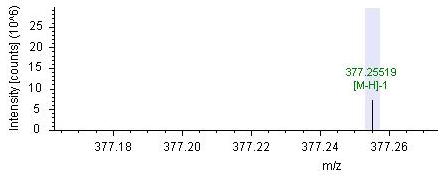

**Figure 3C:** Fragmentation pattern of Gentisic acid leading to signal positive intensities ions [M+H]+1 present in the mass spectrum.

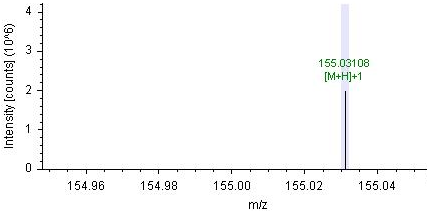

**Figure 3E:** Fragmentation pattern of Glutathione leading to signal intensities ions [M+H]+1 present in the mass spectrum.

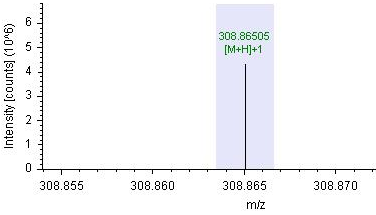

**Figure 3E:** Fragmentation pattern of 4-Hydroxy-butanone leading to signal positive intensities ions [M+H]+1 present in the mass spectrum.

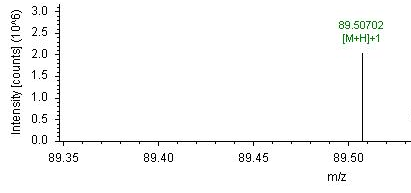
