## Supplemental Table 1 for "Circulating Biomarkers for Therapeutic Response to Immune Checkpoint Inhibitor Therapy in Patients with Advanced Lung Cancer"

**SUPPLEMENTERY TABLES**

**TABLE 1:** The table shows the Raw values of the distribution of expression on the PD-L1, TMB, MSI, and TP53 gene mutations.

| **S. No** | **label** | **TMB** | **MSI (MSI-L=0; MSS= 1)** | **PDL-1 EXPRESSION** | **TP53p** |
| --- | --- | --- | --- | --- | --- |
| 1 | NR | 22.49 | 0 | 0 | 1 |
| 2 | NR | 9.2 | 0 | 0 | 1 |
| 3 | NR | 11.2 | 0 | 0 | 1 |
| 4 | NR | 3.6 | 1 | 0 | 0 |
| 5 | NR | 2.12 | 1 | 0 | 0 |
| 6 | NR | 14.2 | 1 | 0 | 1 |
| 7 | NR | 2.53 | 1 | 0 | 0 |
| 8 | NR | 19.44 | 1 | 0 | 1 |
| 9 | NR | 2.31 | 1 | 0 | 0 |
| 10 | NR | 2.9 | 1 | 0 | 0 |
| 11 | NR | 3.2 | 0 | 0 | 0 |
| 12 | NR | 1.8 | 0 | 0 | 0 |
| 13 | NR | 2.9 | 1 | 0 | 0 |
| 14 | NR | 2.1 | 0 | 0 | 0 |
| 15 | NR | 3.2 | 0 | 0 | 0 |
| 16 | NR | 2.3 | 0 | 0 | 0 |
| 17 | NR | 3.4 | 0 | 0 | 0 |
| 18 | NR | 2.6 | 0 | 0 | 0 |
| 19 | NR | 2.5 | 0 | 0 | 0 |
| 20 | NR | 2.1 | 0 | 0 | 0 |
| 21 | NR | 3.6 | 0 | 0 | 1 |
| 22 | NR | 3.1 | 0 | 0 | 1 |
| 23 | NR | 8.35 | 1 | 0 | 1 |
| 24 | NR | 10.25 | 1 | 0 | 1 |
| 25 | NR | 4.3 | 0 | 0 | 0 |
| 26 | NR | 5.4 | 0 | 0 | 0 |
| 27 | NR | 1.8 | 0 | 0 | 0 |
| 28 | NR | 2.1 | 0 | 0 | 0 |
| 29 | NR | 2.6 | 0 | 0 | 0 |
| 30 | NR | 3.5 | 0 | 0 | 0 |
| 31 | R | 8.35 | 1 | 30 | 0 |
| 32 | R | 6.67 | 1 | 30 | 1 |
| 33 | R | 20.02 | 1 | 60 | 1 |
| 34 | R | 2.1 | 1 | 60 | 1 |
| 35 | R | 10.01 | 0 | 60 | 1 |
| 36 | R | 2.1 | 0 | 60 | 0 |
| 37 | R | 3.4 | 1 | 30 | 0 |
| 38 | R | 3.8 | 1 | 30 | 1 |
| 39 | R | 10.89 | 0 | 30 | 1 |
| 40 | R | 4.16 | 0 | 30 | 0 |
| 41 | R | 14.2 | 1 | 60 | 1 |
| 42 | R | 15.6 | 0 | 30 | 1 |
| 43 | R | 2.51 | 1 | 30 | 0 |
| 44 | R | 10.98 | 1 | 30 | 1 |
| 45 | R | 13.4 | 0 | 30 | 1 |
| 46 | R | 15.6 | 1 | 60 | 1 |
| 47 | R | 14.3 | 1 | 60 | 0 |
| 48 | R | 11.89 | 0 | 60 | 1 |
| 49 | R | 10.7 | 0 | 60 | 1 |
| 50 | R | 20.49 | 1 | 60 | 0 |
| 51 | R | 23.2 | 1 | 60 | 0 |
| 52 | R | 23.35 | 1 | 60 | 1 |
| 53 | R | 20.2 | 1 | 60 | 1 |
| 54 | R | 10.9 | 1 | 60 | 0 |
| 55 | R | 11.5 | 1 | 60 | 0 |
| 56 | R | 17.58 | 0 | 60 | 1 |
| 57 | R | 18.5 | 0 | 60 | 1 |
| 58 | R | 15.6 | 1 | 30 | 0 |
| 59 | R | 17.5 | 1 | 30 | 0 |
| 60 | R | 14.5 | 0 | 30 | 0 |
| 61 | R | 13.6 | 0 | 30 | 0 |
| 62 | R | 13.4 | 0 | 30 | 0 |
| 63 | R | 25.6 | 1 | 60 | 1 |
| 64 | R | 30.5 | 1 | 60 | 1 |
