## Supplemental Table 2 for "Circulating Biomarkers for Therapeutic Response to Immune Checkpoint Inhibitor Therapy in Patients with Advanced Lung Cancer"

**SUPPLEMENTARY TABLE**

**TABLE 2:** The table shows the raw values 576 features which were annotated and identified from a total of 33441 features.

| **kegg Id** | **Molecular Weight** | **RT [min]** | **Derivative** | | | | | | **Validation** | |
| --- | --- | --- | --- | --- | --- | --- | --- | --- | --- | --- |
|  |  |  | **Baseline** | | **8 weeks** | | **16 weeks** | | **baseline** | |
|  |  |  | **NR** | **R** | **NR** | **R** | **NR** | **R** | **NR** | **R** |
| C00003 | 337.04243 | 11.931 | 18688.8 | 12026.14 | 4674.781 | 6767.83 | 11988.92 | 5151.189 | 45558 | 16646.8 |
| C00009 | 254.10487 | 1.709 | 172305 | 99286.09 | 26578.25 | 30983.94 | 34183.77 | 37941.81 | 284398 | 91361.2 |
| C00022 | 99.06808 | 2.626 | 91741.2 | 54369.14 | 10827.01 | 11404.33 | 10321 | 12375.38 | 27219.8 | 92101.9 |
| C00026 | 481.31083 | 14.675 | 66540 | 143573.5 | 36679.5 | 1.17E+05 | 78956.8 | 57116.65 | 30277.1 | 57871.8 |
| C00033 | 1082.6653 | 17.039 | 703242 | 621067.6 | 58430.26 | 62905.98 | 62033.99 | 63656.02 | 787964 | 6.82E+05 |
| C00036 | 1080.6401 | 18.188 | 21929.6 | 26524.99 | 3448.23 | 4258.706 | 3400.123 | 4137.547 | 117032 | 54629.3 |
| C00064 | 444.19877 | 15.926 | 717741 | 294298.6 | 33218.09 | 36685.24 | 39498.5 | 35796.53 | 793789 | 5.61E+05 |
| C00065 | 105.04301 | 1.874 | 43866.5 | 21900.96 | 5480.131 | 5235.887 | 5119.818 | 5016.321 | 32297.7 | 38488.3 |
| C00073 | 145.05242 | 7.504 | 25535.3 | 20648.79 | 5936.984 | 6320.927 | 7347.774 | 6834.231 | 166731 | 1.54E+05 |
| C00082 | 108.02128 | 1.479 | 55559.3 | 57520.94 | 24430.98 | 25413.36 | 21389.73 | 26546.23 | 85978.2 | 45476.3 |
| C00086 | 76.99 | 1.41 | 179261 | 396423.1 | 11348.28 | 11801.12 | 11589.14 | 12285.71 | 35143 | 6.61E+04 |
| C00090 | 561.32605 | 18.364 | 16978.6 | 31194.65 | 5208.158 | 5916.266 | 5037.673 | 6084.675 | 753492 | 6.93E+05 |
| C00097 | 100.5014 | 1.913 | 91263.2 | 50714.41 | 13152.49 | 12283.5 | 11891.25 | 11425.28 | 46804.2 | 57940.2 |
| C00099 | 580.11928 | 15.624 | 122878 | 86499.5 | 25652.76 | 38386.96 | 29964.77 | 27322.87 | 321005 | 64139.5 |
| C00114 | 280.65447 | 20.096 | 25929.5 | 21032.05 | 5307.467 | 6296.003 | 5258.564 | 6293.448 | 146689 | 1.41E+05 |
| C00116 | 198.05141 | 1.504 | 390753 | 389435 | 1.18E+05 | 1.32E+05 | 1.23E+05 | 1.26E+05 | 54857.8 | 55613.9 |
| C00135 | 396.24837 | 13.757 | 107112 | 148267.1 | 71369.1 | 81078.01 | 73920.3 | 82619.38 | 180660 | 93803.8 |
| C00146 | 362.20862 | 11.452 | 58084.6 | 41002.57 | 7342.516 | 8231.818 | 8648.57 | 8044.414 | 147996 | 2.14E+05 |
| C00148 | 392.2917 | 17.052 | 16617.4 | 34502.56 | 4063.012 | 4104.362 | 4470.254 | 3869.003 | 23192.6 | 11680.6 |
| C00153 | 330.27605 | 18.504 | 97248.2 | 90076.84 | 17211.09 | 21609.26 | 17589.82 | 20138.88 | 2.04E+06 | 2.70E+06 |
| C00158 | 279.25575 | 18.464 | 13405.3 | 21500.47 | 4274.629 | 5186.592 | 4054.066 | 4886.217 | 165086 | 1.08E+05 |
| C00159 | 162.11785 | 9.169 | 172046 | 102644.8 | 2.15E+05 | 1.20E+05 | 1.90E+05 | 1.42E+05 | 408276 | 2.06E+05 |
| C00163 | 177.05023 | 11.536 | 84006.1 | 79515.33 | 13727.77 | 15254.62 | 16040.34 | 15744.01 | 6.47E+06 | 6.86E+06 |
| C00186 | 237.82322 | 1.629 | 26379.9 | 27734.66 | 2437.999 | 2586.052 | 2562.223 | 2381.53 | 382412 | 1.30E+05 |
| C00196 | 592.35985 | 14.605 | 252633 | 493698.4 | 1.87E+05 | 3.39E+05 | 2.46E+05 | 2.08E+05 | 14365.5 | 24435.6 |
| C00219 | 335.28795 | 10.227 | 73708 | 35990.53 | 8889.7 | 9271.12 | 9419.273 | 8904.46 | 178365 | 1.99E+05 |
| C00224 | 674.2391 | 1.583 | 130512 | 72195.05 | 31432.93 | 28761.81 | 31788.16 | 37137.18 | 6381.89 | 6686.74 |
| C00230 | 100.05153 | 3.56 | 87113.3 | 83228.09 | 34112.81 | 26493.31 | 2.03E+05 | 1.02E+05 | 43161.4 | 29164.3 |
| C00232 | 200.02287 | 11.091 | 6.31E+07 | 4.32E+07 | 6.23E+07 | 3.83E+07 | 5.06E+07 | 4.39E+07 | 209436 | 67822.4 |
| C00239 | 394.23497 | 18.681 | 17677.2 | 18253.07 | 3424.239 | 4597 | 3652.377 | 4104.761 | 38453.6 | 36235.5 |
| C00249 | 140.04699 | 7.757 | 515318 | 454440.6 | 1.20E+05 | 1.17E+05 | 1.18E+05 | 1.30E+05 | 1.31E+06 | 1.02E+06 |
| C00251 | 280.6543 | 17.181 | 264324 | 267029.2 | 1.47E+05 | 2.19E+05 | 3.23E+05 | 2.19E+05 | 589386 | 5.68E+05 |
| C00253 | 332.23403 | 19.616 | 39436.2 | 39479.76 | 10523.78 | 11872.22 | 10128.73 | 11681.09 | 754610 | 5.19E+05 |
| C00261 | 354.13582 | 2.184 | 353741 | 182129.3 | 1.11E+05 | 1.62E+05 | 1.40E+05 | 1.03E+05 | 295335 | 3.29E+05 |
| C00262 | 778.8465 | 17.441 | 103592 | 69682.73 | 6693.621 | 7863.527 | 6744.692 | 7576.296 | 65079.6 | 34976 |
| C00270 | 180.07601 | 3.897 | 6381.46 | 10008.05 | 8820.447 | 10106.93 | 11280.94 | 7731.849 | 61018.4 | 18412 |
| C00279 | 406.69759 | 10.177 | 24561.6 | 14192.7 | 21937.02 | 27689.6 | 13007.42 | 9437.749 | 223586 | 62809.6 |
| C00292 | 558.32645 | 14.457 | 196700 | 244288.1 | 1.59E+05 | 2.05E+05 | 2.73E+05 | 1.53E+05 | 13735.6 | 11855.2 |
| C00295 | 138.06778 | 14.56 | 106342 | 59144.23 | 12438.54 | 14675.59 | 12874.77 | 13183.68 | 164499 | 1.48E+05 |
| C00296 | 237.9821 | 21.501 | 277440 | 310204.5 | 1.26E+05 | 1.24E+05 | 1.08E+05 | 1.13E+05 | 336567 | 4.07E+05 |
| C00300 | 112.0526 | 14.243 | 26551.5 | 19029.38 | 4358.434 | 5332.795 | 5398.243 | 5128.823 | 97645.6 | 98563.8 |
| C00302 | 200.02277 | 1.82 | 140792 | 70683.7 | 14468.69 | 15535.56 | 14474.48 | 15088.66 | 342092 | 2.91E+05 |
| C00307 | 266.18764 | 10.847 | 60636.5 | 36829.24 | 8418.708 | 9609.798 | 9028.361 | 9060.677 | 42878.9 | 57998.2 |
| C00315 | 118.06216 | 3.066 | 2.27E+06 | 2.34E+06 | 1.38E+06 | 1.64E+06 | 1.21E+06 | 1.20E+06 | 224261 | 1.40E+05 |
| C00327 | 330.23984 | 12.499 | 38861.3 | 34596.96 | 13700.04 | 19061.5 | 17892.91 | 14470.15 | 299905 | 5.03E+05 |
| C00328 | 145.05247 | 3.967 | 37100.7 | 50045.62 | 10101.05 | 11924.09 | 11313.52 | 12121.12 | 2.56E+06 | 1.56E+06 |
| C00331 | 410.22792 | 13.821 | 69344.6 | 51806.93 | 13891.6 | 14903.85 | 17649.21 | 16017.04 | 219908 | 5.39E+05 |
| C00334 | 281.92552 | 22.377 | 693292 | 1.15E+06 | 7.18E+05 | 5.95E+05 | 7.36E+05 | 7.55E+05 | 163803 | 1.48E+05 |
| C00352 | 381.26361 | 10.559 | 116745 | 83388.76 | 28754.1 | 31359.96 | 28008.98 | 34044.71 | 96855.4 | 14084.6 |
| C00353 | 580.13807 | 17.794 | 27478 | 22441.86 | 2852.502 | 3390.894 | 2773.034 | 3324.68 | 15736.3 | 54450.4 |
| C00364 | 335.06799 | 3.456 | 17727.3 | 21087.52 | 10768.69 | 13382.17 | 3862.334 | 4923.463 | 47067.8 | 35484.7 |
| C00366 | 298.25021 | 17.225 | 25256.2 | 19244.97 | 8869.801 | 8814.955 | 7140.427 | 9330.246 | 150137 | 4.20E+05 |
| C00407 | 425.22457 | 10.505 | 1.23E+07 | 6.73E+06 | 5.26E+06 | 2.90E+06 | 3.91E+06 | 4.11E+06 | 174646 | 65659 |
| C00417 | 279.64696 | 17.554 | 53682.5 | 45530.36 | 4598.62 | 5364.003 | 4579.979 | 5230.697 | 21722.7 | 54525.1 |
| C00440 | 605.33986 | 9.226 | 45865.5 | 28311.08 | 3032.351 | 3421.756 | 3161.108 | 3311.168 | 32865.7 | 4974.43 |
| C00460 | 182.09416 | 2.037 | 9896.17 | 10079.5 | 4261.11 | 5556.1 | 5395.016 | 3574.294 | 31907.2 | 25691.9 |
| C00482 | 148.0661 | 4.01 | 248718 | 433797 | 1.29E+05 | 5.05E+05 | 5.43E+05 | 3.50E+05 | 710741 | 3.30E+05 |
| C00486 | 578.14164 | 22.881 | 25825.8 | 29968.37 | 1.49E+05 | 1.06E+05 | 1.91E+05 | 51901.75 | 64473.3 | 47583.4 |
| C00534 | 85.05238 | 2.051 | 20917.5 | 16685.38 | 8181.946 | 9571.437 | 9899.715 | 8500.133 | 33636.4 | 36585.7 |
| C00550 | 156.04209 | 7.814 | 756506 | 259058.5 | 54259.59 | 52453.58 | 53315.96 | 55156.29 | 921097 | 6.23E+05 |
| C00559 | 323.15149 | 11.195 | 17672.6 | 5886.57 | 10654.13 | 7474.632 | 13626.47 | 8421.035 | 216040 | 39501.7 |
| C00590 | 406.11026 | 12.505 | 103824 | 133026 | 15680.29 | 17373.61 | 16560.5 | 16300.65 | 458634 | 3.62E+05 |
| C00596 | 541.33207 | 14.356 | 679996 | 1.48E+06 | 4.54E+05 | 6.77E+05 | 4.00E+05 | 5.76E+05 | 12875 | 8464.12 |
| C00612 | 336.22808 | 10.827 | 13149.7 | 10979.92 | 15162.93 | 30086.22 | 16493.34 | 13176.24 | 233134 | 30464.7 |
| C00628 | 154.02385 | 1.701 | 571524 | 189156.9 | 4276.726 | 4829.848 | 4312.424 | 4724.22 | 5934274 | 2785462 |
| C00633 | 318.27617 | 17.314 | 221518 | 266954 | 34271.58 | 40017.28 | 42685.45 | 39515.32 | 7.66E+06 | 6.15E+06 |
| C00644 | 467.30044 | 10.611 | 20921.8 | 15523.69 | 5913.337 | 17669.08 | 7750.81 | 9143.687 | 12426.8 | 52993.8 |
| C00651 | 337.20911 | 14.852 | 78355.4 | 41862.81 | 10703.44 | 12057.03 | 13252.06 | 11670.17 | 624152 | 1.54E+05 |
| C00666 | 657.31444 | 13.996 | 24571.9 | 25921.39 | 10353.74 | 32120.02 | 34006.81 | 22320.77 | 22903.9 | 31763.1 |
| C00669 | 238.08817 | 12.358 | 103424 | 37776.19 | 3226.984 | 3350.941 | 3351.764 | 3234.227 | 7585.66 | 15548 |
| C00670 | 118.06151 | 3.331 | 163047 | 81946.53 | 34266.82 | 57396.15 | 41117.07 | 44180.43 | 5.52E+06 | 5.65E+06 |
| C00695 | 280.92496 | 17.799 | 112285 | 91332.49 | 8817.44 | 10510.1 | 8557.01 | 10495.79 | 98247 | 36191.5 |
| C00712 | 152.11981 | 12.43 | 77439.7 | 107829.5 | 9015.146 | 9808.485 | 9692.704 | 9699.365 | 355705 | 2.49E+05 |
| C00735 | 153.99807 | 21.194 | 748012 | 867868.6 | 6.20E+05 | 5.85E+05 | 5.31E+05 | 4.48E+05 | 261353 | 1.65E+05 |
| C00740 | 128.05222 | 18.695 | 24739.4 | 43873.44 | 10135.24 | 13952.18 | 9865.95 | 10936.76 | 55225 | 27573.5 |
| C00760 | 430.30721 | 7.86 | 169465 | 204807.3 | 1.12E+05 | 1.11E+05 | 4.71E+05 | 2.99E+05 | 162716 | 86486.1 |
| C00761 | 407.28455 | 12.523 | 60743.5 | 32405.84 | 6470.947 | 6794.656 | 8074.568 | 6980.226 | 333917 | 88036.2 |
| C00762 | 323.16902 | 9.528 | 545685 | 426799.4 | 49688.2 | 62971.3 | 46076.94 | 61120.15 | 1.17E+07 | 5.22E+06 |
| C00763 | 599.27193 | 2.225 | 408436 | 252040.9 | 2.18E+05 | 1.89E+05 | 2.69E+05 | 1.79E+05 | 97534.6 | 1.00E+05 |
| C00785 | 154.02806 | 1.545 | 188465 | 87485.39 | 39480.86 | 41688.05 | 37833.92 | 40729.76 | 15297.4 | 14471.2 |
| C00791 | 902.55569 | 16.882 | 166799 | 89058.04 | 7430.87 | 8693.006 | 7801.708 | 8331.372 | 318924 | 2.91E+05 |
| C00792 | 218.07897 | 7.104 | 543290 | 484633.6 | 2.64E+05 | 5.10E+05 | 6.80E+05 | 1.20E+06 | 566492 | 1.97E+05 |
| C00802 | 194.07262 | 2.035 | 48593.7 | 38590.32 | 19075.72 | 23279.41 | 23531.61 | 18756.78 | 447412 | 6.78E+05 |
| C00819 | 162.12526 | 9.435 | 150209 | 143102.9 | 20574.92 | 21742.5 | 22504.23 | 20880.07 | 810292 | 1.16E+06 |
| C00828 | 499.18344 | 2.061 | 35297.6 | 12864.67 | 10890.52 | 17610.08 | 33945.8 | 18654.12 | 34650.1 | 44778.3 |
| C00835 | 364.25167 | 8.915 | 32089.2 | 22705.64 | 2402.851 | 2615.948 | 2643.075 | 2618.41 | 22234.6 | 8926.21 |
| C00836 | 118.06149 | 3.088 | 78919.5 | 46981.87 | 12974.92 | 16479.43 | 17383.87 | 13012.23 | 1.54E+06 | 1.74E+06 |
| C00864 | 114.04263 | 1.856 | 522921 | 278713.7 | 57810.8 | 59987.87 | 55749.21 | 57415.93 | 2.81E+06 | 2.51E+06 |
| C00870 | 532.30532 | 14.937 | 272764 | 261518.1 | 1.06E+06 | 1.64E+06 | 2.15E+06 | 1.35E+06 | 46961.7 | 11999.3 |
| C00881 | 408.11891 | 10.866 | 46678.8 | 25431.42 | 6846.617 | 8269.072 | 7098.677 | 6570.979 | 103791 | 1.33E+05 |
| C00882 | 248.07746 | 7.378 | 14045.4 | 10378.96 | 3585.453 | 3797.497 | 4396.635 | 4152.389 | 98468.3 | 14961.4 |
| C00902 | 168.11474 | 16.569 | 927642 | 303120.3 | 21778.26 | 24201.63 | 22762.96 | 24464.55 | 283499 | 1.48E+05 |
| C00906 | 147.08845 | 1.744 | 277071 | 171525.7 | 56112.99 | 59704.64 | 54494.92 | 61541.95 | 236271 | 1.98E+05 |
| C00922 | 592.46617 | 14.48 | 44101.1 | 57433.95 | 11152.68 | 24310.5 | 1.85E+05 | 20774.01 | 21989 | 18956.4 |
| C00931 | 237.13593 | 14.937 | 32187.8 | 17470.17 | 4583.968 | 4743.808 | 5334.821 | 4684.748 | 32755.6 | 26727.7 |
| C00940 | 126.01911 | 16.528 | 313549 | 107491 | 8356.978 | 9293.024 | 8861.481 | 9418.185 | 107898 | 93186.3 |
| C00951 | 578.14177 | 15.496 | 102626 | 118931.4 | 75019.54 | 51744.72 | 69626.88 | 83121.71 | 1.13E+06 | 1.75E+05 |
| C00990 | 336.19009 | 10.603 | 58522.6 | 80366.83 | 64532.76 | 80223.19 | 4.51E+05 | 44199.32 | 319587 | 52487.4 |
| C01004 | 1023.6081 | 12.695 | 19583.6 | 31621.17 | 3032.384 | 3379.373 | 3071.273 | 3225.188 | 517585 | 1.98E+05 |
| C01013 | 541.33188 | 14.444 | 56641.2 | 34937.57 | 4392.677 | 4232.349 | 5157.952 | 4090.878 | 11521 | 17791.1 |
| C01047 | 143.09437 | 2.969 | 218920 | 169744.5 | 27403.46 | 28795.18 | 28877.39 | 30106.47 | 1.92E+06 | 1.74E+06 |
| C01053 | 330.23996 | 13.178 | 153604 | 81319.5 | 22231.55 | 22958.68 | 23688.78 | 20125.51 | 390638 | 3.87E+05 |
| C01089 | 226.16739 | 6.875 | 411591 | 284490.1 | 96123.59 | 1.30E+05 | 86452.03 | 91545.36 | 455161 | 1.58E+05 |
| C01091 | 156.88039 | 2.02 | 44261.4 | 42468.4 | 12191.18 | 19603.7 | 19518.28 | 13403.83 | 23500.7 | 19401 |
| C01092 | 338.24342 | 19.912 | 35210 | 29929.71 | 7707.703 | 9062.65 | 8446.059 | 9785.041 | 852395 | 3.07E+06 |
| C01100 | 346.24983 | 12.153 | 101006 | 45101.43 | 4367.799 | 4792.495 | 4764.345 | 4500.443 | 54846.7 | 50141.3 |
| C01109 | 337.12081 | 14.202 | 41057.8 | 27466.02 | 6532.757 | 8094.418 | 8832.999 | 7692.773 | 1.85E+06 | 2.28E+06 |
| C01120 | 449.80367 | 11.31 | 14022.7 | 4577.227 | 9292.457 | 11015.37 | 12209.84 | 7748.672 | 42664.8 | 17528.8 |
| C01142 | 168.11464 | 8.914 | 29449.1 | 37526.99 | 3787.559 | 4055.797 | 4494.536 | 4000.02 | 103212 | 1.09E+05 |
| C01147 | 595.12855 | 14.492 | 47991.2 | 29901.68 | 21738.41 | 29054.62 | 23318.71 | 32054.3 | 17586.3 | 22056.8 |
| C01152 | 167.04725 | 2.033 | 83951.5 | 55213.01 | 28031.15 | 26051.73 | 27028.38 | 18636.49 | 77531.9 | 90335.3 |
| C01169 | 306.04774 | 10.752 | 25345.8 | 26427.33 | 10055.76 | 11210.74 | 12677.52 | 10464.46 | 43354.5 | 35343.3 |
| C01173 | 623.35066 | 17.071 | 340557 | 221074.6 | 19524.77 | 22045.09 | 19297.85 | 21528.94 | 636189 | 6.69E+05 |
| C01180 | 129.00241 | 1.458 | 87793.9 | 131821.3 | 26623.78 | 32637.39 | 29701.7 | 29471.08 | 269950 | 96418.5 |
| C01187 | 249.11073 | 8.19 | 59182.3 | 46081.9 | 23813.43 | 23758.5 | 24242.29 | 14512.5 | 396526 | 1.34E+05 |
| C01235 | 480.23494 | 18.767 | 53179.8 | 58401.25 | 11334.85 | 14198.84 | 12119.19 | 14123.86 | 1.29E+06 | 4.73E+05 |
| C01250 | 180.06352 | 3.434 | 207321 | 302522.4 | 1.79E+05 | 1.28E+05 | 1.38E+05 | 1.46E+05 | 77351 | 54665.6 |
| C01267 | 176.03958 | 11.741 | 125412 | 135509.2 | 52109.54 | 43399 | 67516.22 | 1.02E+05 | 184348 | 60974.4 |
| C01300 | 542.33553 | 14.848 | 743356 | 728401.1 | 4.66E+06 | 5.26E+06 | 1.03E+07 | 4.79E+06 | 42145.9 | 35739.7 |
| C01301 | 473.94949 | 16.136 | 18166.2 | 6886.029 | 4560.235 | 5240.657 | 6307.405 | 4315.41 | 19844.6 | 20311.9 |
| C01403 | 335.28806 | 10.135 | 18271.9 | 19148.55 | 5397.876 | 12246.45 | 11308.65 | 8506.212 | 242757 | 35288.9 |
| C01411 | 154.13541 | 14.213 | 99075.6 | 69206.33 | 15918.3 | 19754.52 | 21165.96 | 18758.02 | 285894 | 2.16E+05 |
| C01432 | 330.23985 | 13.509 | 301821 | 244001.9 | 29014.48 | 31841.1 | 31769.18 | 30911.33 | 4.43E+06 | 1.80E+06 |
| C01479 | 779.84368 | 16.344 | 161690 | 100399.1 | 9429.979 | 10291.72 | 9638.528 | 10150.25 | 481778 | 6.42E+05 |
| C01493 | 541.33259 | 15.804 | 228810 | 621816.8 | 2.62E+05 | 4.53E+05 | 4.91E+05 | 9.75E+05 | 84424.1 | 72084.5 |
| C01494 | 781.7926 | 3.289 | 2.51E+06 | 712715.5 | 3.97E+05 | 4.86E+05 | 3.92E+05 | 2.57E+06 | 70307.8 | 70118.6 |
| C01507 | 451.93291 | 17.179 | 207252 | 101271.7 | 9394.501 | 10794.41 | 9241.311 | 10392.82 | 144808 | 1.45E+05 |
| C01530 | 129.04301 | 13.704 | 109902 | 121681.6 | 14957.26 | 17091.06 | 16109.09 | 17114.05 | 3.49E+06 | 4.21E+06 |
| C01561 | 451.27784 | 10.695 | 6546.75 | 6849.458 | 3522.701 | 17190.88 | 8427.684 | 10374.68 | 34629.9 | 24481.9 |
| C01586 | 810.41726 | 2.506 | 5.77E+06 | 3.34E+06 | 2.07E+06 | 1.74E+06 | 1.54E+06 | 1.81E+06 | 47705.4 | 1.05E+05 |
| C01591 | 391.28612 | 16.682 | 50378.1 | 36245.02 | 4214.426 | 4180.487 | 4848.556 | 3899.973 | 14792 | 10049.5 |
| C01595 | 425.28914 | 9.882 | 2.09E+07 | 1.40E+07 | 3.34E+07 | 2.86E+07 | 4.22E+07 | 2.29E+07 | 872651 | 2.16E+05 |
| C01606 | 135.06822 | 1.882 | 173805 | 90620.28 | 21965.74 | 22571.95 | 21344.35 | 20976.2 | 2.00E+06 | 2.09E+06 |
| C01613 | 342.11542 | 14.707 | 18771.1 | 7229.261 | 4870.592 | 6664.431 | 4478.828 | 2892.813 | 13473.1 | 3888.17 |
| C01620 | 108.02108 | 1.083 | 201662 | 176212.5 | 77673.84 | 1.08E+05 | 88862.53 | 84669.26 | 558276 | 5.45E+05 |
| C01672 | 753.35156 | 2.513 | 97317.7 | 83481.04 | 42010.5 | 52777.38 | 1.16E+05 | 48694.59 | 442711 | 5.42E+05 |
| C01673 | 452.14019 | 12.466 | 3053.01 | 5497.238 | 7954.028 | 3349.737 | 3511.473 | 3315.859 | 372284 | 36056.2 |
| C01674 | 568.32364 | 18.107 | 31967.7 | 27025.79 | 4616.747 | 5690.645 | 4562.225 | 5868.448 | 83390.7 | 51018.5 |
| C01690 | 350.9219 | 7.993 | 61308 | 31535.76 | 11195.59 | 10533.93 | 10848.23 | 9589.383 | 2.10E+06 | 4.38E+06 |
| C01746 | 216.17153 | 2.463 | 3.51E+06 | 1.67E+06 | 5.27E+06 | 2.40E+06 | 5.37E+06 | 4.06E+06 | 1.64E+06 | 3.87E+06 |
| C01747 | 99.06856 | 2.63 | 572680 | 367857.2 | 70797.52 | 74453.95 | 68074.07 | 80874.23 | 4.35E+06 | 4.54E+06 |
| C01772 | 542.33565 | 15.32 | 267177 | 413596.6 | 4.60E+05 | 7.81E+05 | 8.15E+05 | 8.83E+05 | 136257 | 38307.9 |
| C01792 | 569.36516 | 20.273 | 32704.3 | 34566.35 | 8100.241 | 9132.769 | 8005.884 | 9282.123 | 58616.1 | 1.03E+05 |
| C01826 | 132.08967 | 1.513 | 202217 | 63245.18 | 18223.7 | 19006.37 | 19279.42 | 19202.27 | 1.35E+06 | 8.93E+05 |
| C01909 | 406.12895 | 10.852 | 34268.3 | 24516.75 | 52335.86 | 76852.52 | 1.47E+05 | 70374.51 | 643377 | 1.27E+05 |
| C01921 | 198.02827 | 1.482 | 1.90E+06 | 1.05E+06 | 5.98E+05 | 5.34E+05 | 4.84E+05 | 4.65E+05 | 230201 | 92228.4 |
| C01924 | 845.59154 | 2.336 | 1.79E+06 | 1.22E+06 | 4.05E+05 | 4.03E+05 | 3.18E+05 | 3.78E+05 | 415792 | 2.08E+05 |
| C01962 | 133.97319 | 1.663 | 90931.1 | 56235 | 12861.15 | 13273.48 | 13887.52 | 13702.82 | 63245.1 | 62477.5 |
| C02076 | 95.03736 | 12.996 | 44600.2 | 53892.66 | 4615.648 | 5075.315 | 5654.152 | 4857.233 | 416214 | 3.81E+05 |
| C02115 | 486.38807 | 14.45 | 57283.5 | 34997.07 | 4158.702 | 4346.983 | 5286.688 | 4215.309 | 45040.1 | 2.32E+05 |
| C02117 | 456.16933 | 14.147 | 60158.6 | 35030.67 | 6208.63 | 6536.029 | 7581.505 | 6083.395 | 14607.8 | 18611.9 |
| C02126 | 337.11937 | 1.952 | 1.06E+06 | 667375.2 | 3.01E+05 | 3.25E+05 | 3.19E+05 | 4.34E+05 | 161045 | 2.54E+05 |
| C02165 | 351.2301 | 11.335 | 14081.8 | 18075.55 | 4879.691 | 5432.285 | 4911.199 | 4703.27 | 6434.42 | 12250.9 |
| C02214 | 226.0055 | 10.8 | 31119.8 | 22498.23 | 9012.884 | 10176.39 | 11640.02 | 8518.238 | 26082.8 | 6003.13 |
| C02230 | 568.32093 | 17.096 | 154300 | 93508.65 | 8985.253 | 9758.115 | 8537.257 | 9716.389 | 86042.1 | 69094.1 |
| C02240 | 335.31796 | 14.448 | 846922 | 306323.7 | 1.34E+05 | 2.17E+05 | 2.44E+05 | 2.01E+05 | 116662 | 1.28E+05 |
| C02289 | 150.10415 | 10.086 | 151827 | 69991.92 | 17435.87 | 18418.79 | 18657.31 | 18598.42 | 257692 | 3.41E+05 |
| C02351 | 444.844 | 16.947 | 37745.5 | 42369.49 | 4281.129 | 4709.715 | 4459.884 | 4744.525 | 81413.4 | 1.41E+05 |
| C02362 | 146.00997 | 1.46 | 18293.4 | 22070.02 | 7775.801 | 5889.996 | 5406.925 | 5546.409 | 190216 | 73450.7 |
| C02367 | 403.98328 | 9.841 | 1.71E+06 | 823159.1 | 3.68E+05 | 3.77E+05 | 3.56E+05 | 3.65E+05 | 26067.2 | 20889.4 |
| C02372 | 406.04173 | 10.579 | 212557 | 238201.4 | 1.13E+05 | 1.24E+05 | 1.64E+05 | 1.02E+05 | 66658.8 | 26505.4 |
| C02376 | 335.31819 | 15.055 | 2.26E+06 | 707564.1 | 1.52E+06 | 1.34E+06 | 1.49E+06 | 1.44E+06 | 225822 | 2.59E+05 |
| C02395 | 639.61293 | 9.387 | 9157.28 | 15070.94 | 5559.964 | 5913.012 | 5070.877 | 5306.57 | 106183 | 19612.4 |
| C02486 | 471.09392 | 10.544 | 41827.1 | 6840.904 | 4854.799 | 9185.996 | 53249.68 | 7276.166 | 59506.9 | 6122 |
| C02502 | 540.32022 | 14.889 | 75944.4 | 59771.88 | 1.68E+05 | 2.80E+05 | 4.36E+05 | 1.79E+05 | 97747.9 | 51854 |
| C02505 | 582.32405 | 14.45 | 38262.6 | 67496.55 | 49318.07 | 69997.87 | 39056.8 | 70255.87 | 11846.9 | 12194.2 |
| C02528 | 562.31266 | 20.524 | 89457.8 | 85900.03 | 16889.96 | 18281.16 | 18396.12 | 18709.05 | 257947 | 3.53E+05 |
| C02535 | 124.95232 | 1.455 | 19111.1 | 14940.03 | 3578.471 | 3693.278 | 2581.183 | 4623.758 | 104160 | 89466.5 |
| C02571 | 279.16362 | 13.457 | 24672.9 | 24438.28 | 12126.14 | 10917.08 | 14438.31 | 12965.04 | 93704.7 | 87691.1 |
| C02648 | 417.03529 | 9.697 | 5725.89 | 3989.263 | 7389.071 | 47312.35 | 77101.36 | 30194.15 | 40610.3 | 26101.5 |
| C02659 | 318.27608 | 17.577 | 2.41E+06 | 1.86E+06 | 1.40E+06 | 1.15E+06 | 1.52E+06 | 1.24E+06 | 2.32E+06 | 1.35E+06 |
| C02700 | 402.16517 | 19.926 | 37639.6 | 30401.06 | 7869.298 | 9215.389 | 8759.059 | 9777.106 | 67628.1 | 65611.5 |
| C02771 | 606.32374 | 17.248 | 362035 | 206261.4 | 18740.17 | 21673.34 | 18764.38 | 21510.29 | 617576 | 7.55E+05 |
| C02777 | 649.78032 | 14.063 | 39217.4 | 61613.29 | 69033.33 | 2.41E+05 | 2.69E+05 | 1.05E+05 | 17214.3 | 30973.7 |
| C02838 | 108.02116 | 7.741 | 566219 | 397627.3 | 93299.48 | 99895.52 | 1.01E+05 | 1.05E+05 | 4.53E+06 | 2.99E+06 |
| C02862 | 754.56211 | 2.758 | 28159.6 | 28537.9 | 56039.08 | 58809.98 | 94903.8 | 48095.15 | 375182 | 3.09E+05 |
| C02876 | 487.17097 | 14.994 | 88960 | 70881.16 | 62774.97 | 59076.12 | 81371.69 | 65245.41 | 103472 | 17136.9 |
| C02938 | 329.16859 | 10.01 | 1.00E+06 | 666759.9 | 3.62E+06 | 2.29E+06 | 4.10E+06 | 3.07E+06 | 13595.2 | 9864.36 |
| C02982 | 237.84408 | 2.027 | 123746 | 78335.13 | 33558.6 | 35430.94 | 32169.65 | 39785.88 | 59984 | 93913.4 |
| C02990 | 132.08993 | 1.882 | 348973 | 188979.5 | 43790.7 | 43010.56 | 40503.5 | 39431.4 | 6.24E+06 | 6.24E+06 |
| C03001 | 525.74862 | 17.68 | 374965 | 143296 | 14636.07 | 17337.45 | 15014.75 | 17267.56 | 4.22E+06 | 3.96E+06 |
| C03017 | 234.16121 | 1.909 | 253739 | 93199.5 | 33901.86 | 32013.26 | 31620.05 | 35478.52 | 172088 | 1.65E+05 |
| C03028 | 168.01852 | 1.838 | 101432 | 153433.3 | 88903.82 | 49579.79 | 48358.73 | 93067.56 | 37862.9 | 38143.3 |
| C03033 | 989.48551 | 17.074 | 67668.5 | 54308.13 | 25525.31 | 63704.39 | 23154.19 | 19468.45 | 36606.8 | 52719.7 |
| C03113 | 220.09958 | 7.758 | 496499 | 382454.8 | 88156.69 | 84640.57 | 85366.07 | 92636.62 | 343928 | 2.67E+05 |
| C03137 | 143.09442 | 1.999 | 42126.2 | 66782.6 | 27865.14 | 24347.91 | 11455.93 | 18352.7 | 458329 | 3.16E+05 |
| C03150 | 148.05236 | 2.349 | 209028 | 206213.7 | 39696.08 | 44424.12 | 38378.34 | 44878.57 | 159827 | 7.49E+05 |
| C03174 | 198.04088 | 1.702 | 3.14E+06 | 2.17E+06 | 4.93E+05 | 1.02E+06 | 9.01E+05 | 6.22E+05 | 498657 | 9.99E+05 |
| C03199 | 558.32744 | 17.038 | 89901 | 74785.6 | 6655.325 | 7579.237 | 6641.935 | 7437.587 | 25697 | 46397.3 |
| C03205 | 406.17011 | 10.029 | 270230 | 232968.9 | 1.50E+05 | 1.46E+05 | 1.88E+05 | 1.22E+05 | 461904 | 73830.7 |
| C03215 | 555.82994 | 22.138 | 19206.4 | 10304.27 | 11291.65 | 11349.21 | 11422.88 | 11542.7 | 128608 | 1.57E+05 |
| C03219 | 487.235 | 14.691 | 228265 | 86637.62 | 51912.85 | 57236.51 | 78667.01 | 43525.31 | 29901.5 | 47553.9 |
| C03228 | 362.20851 | 9.975 | 27758.1 | 12602.14 | 3521.276 | 3568.233 | 3585.768 | 3318.302 | 6475 | 16606.4 |
| C03232 | 404.18052 | 10.909 | 491416 | 270842.7 | 1.87E+05 | 1.55E+05 | 1.98E+05 | 1.81E+05 | 112988 | 71217.7 |
| C03242 | 337.33391 | 18.524 | 32064 | 46454.56 | 9653.337 | 11977.76 | 9621.242 | 11219.75 | 638356 | 4.70E+05 |
| C03338 | 585.7236 | 14.686 | 14702.5 | 13518.16 | 27879.22 | 72817.81 | 45298.8 | 34444.69 | 24813.5 | 38715.1 |
| C03366 | 256.96939 | 14.377 | 34856.3 | 27351.14 | 5741.904 | 7216.136 | 6328.551 | 5723.95 | 14540.3 | 15492.7 |
| C03404 | 401.181 | 17.874 | 139461 | 109776.8 | 12857.31 | 16171.05 | 13237.05 | 15290.02 | 91136.9 | 3.17E+05 |
| C03448 | 150.10413 | 10.877 | 120712 | 63133.81 | 16938.26 | 20573.61 | 17314.58 | 16196.23 | 265220 | 1.42E+05 |
| C03510 | 453.28255 | 12.598 | 60859.2 | 48438.8 | 2.07E+05 | 1.46E+05 | 1.73E+05 | 2.51E+05 | 388367 | 4.54E+05 |
| C03621 | 143.09522 | 12.67 | 44073.4 | 38300.51 | 3957.162 | 4571.831 | 4079.93 | 4263.623 | 110507 | 3.32E+05 |
| C03664 | 532.30468 | 14.602 | 2.50E+06 | 4.88E+06 | 9.35E+05 | 1.78E+06 | 8.61E+05 | 1.11E+06 | 18864.4 | 20713.6 |
| C03672 | 330.23988 | 14.234 | 407160 | 228846.4 | 48648.69 | 59969.9 | 62607.53 | 55815.96 | 2.33E+06 | 1.45E+06 |
| C03684 | 320.12854 | 10.624 | 50353 | 32844.5 | 9536.626 | 9656.842 | 9488.637 | 10695.29 | 107306 | 52219.6 |
| C03837 | 337.11949 | 1.575 | 57975.2 | 47585.52 | 77033.93 | 74246.24 | 78942.02 | 1.01E+05 | 6420.48 | 4901.77 |
| C03846 | 698.74672 | 14.314 | 78850.5 | 60918.9 | 8096.755 | 9759.414 | 9178.243 | 9115.112 | 1.82E+06 | 3.02E+06 |
| C03852 | 400.27935 | 14.763 | 30176.6 | 32720.67 | 3469.517 | 3852.517 | 4504.724 | 3746.105 | 12760.8 | 6014.05 |
| C03887 | 195.10442 | 7.347 | 29592.7 | 17645.59 | 27826.54 | 1.76E+05 | 6.18E+05 | 1.77E+05 | 123253 | 77754.1 |
| C03901 | 119.05794 | 1.676 | 15095.6 | 8603.016 | 17574.21 | 5831.468 | 5798.375 | 9224.32 | 18377.9 | 22038.3 |
| C03916 | 1080.6401 | 17.034 | 134784 | 90058.64 | 8453.803 | 9639.974 | 8967.043 | 9489.68 | 46840.4 | 80818 |
| C03943 | 166.09901 | 8.776 | 112095 | 105406.5 | 8746.187 | 9367.44 | 9175.378 | 9332.572 | 197668 | 1.18E+05 |
| C03962 | 150.10414 | 10.409 | 190545 | 225726.6 | 59753.97 | 66133.96 | 64700.45 | 68897.66 | 295863 | 2.77E+05 |
| C04036 | 541.3319 | 18.275 | 13750.3 | 17941.61 | 3154.834 | 3847.072 | 2950.789 | 4104.26 | 208763 | 1.97E+05 |
| C04067 | 519.33126 | 14.886 | 2.98E+06 | 5.91E+06 | 2.05E+06 | 3.22E+06 | 1.72E+06 | 1.88E+06 | 60491.2 | 45565.4 |
| C04084 | 222.19798 | 14.155 | 75265.2 | 53315.12 | 11494.74 | 13958.41 | 15868.5 | 13534.82 | 1.48E+06 | 1.98E+06 |
| C04100 | 475.23701 | 17.047 | 481417 | 424692.3 | 30400.15 | 33507.64 | 31189.51 | 32853.6 | 952631 | 6.91E+05 |
| C04102 | 541.33198 | 18.457 | 55743.5 | 77490.17 | 14683.31 | 16945.7 | 13043.36 | 16359.3 | 453924 | 2.94E+05 |
| C04103 | 239.15363 | 19.769 | 22687.9 | 17710.01 | 4257.344 | 5497.32 | 4506.553 | 5319.866 | 198625 | 1.18E+05 |
| C04104 | 367.07927 | 11.905 | 1.06E+08 | 6.95E+07 | 2.49E+07 | 2.50E+07 | 2.87E+07 | 2.38E+07 | 406060 | 54975.8 |
| C04110 | 149.01914 | 13.592 | 134361 | 138008.7 | 17338.85 | 19859.36 | 18727.37 | 19726.81 | 2.55E+07 | 6.26E+07 |
| C04122 | 82.04939 | 19.806 | 54384.6 | 65227.25 | 15607.58 | 22561.01 | 10343.85 | 26949.16 | 65081.4 | 41073.6 |
| C04148 | 148.05216 | 2.746 | 191530 | 187647.5 | 29210.99 | 33834.72 | 28944.97 | 36343.58 | 2.58E+08 | 1.23E+08 |
| C04277 | 238.08136 | 22.233 | 6.37E+06 | 6.39E+06 | 2.00E+06 | 2.15E+06 | 1.34E+06 | 1.83E+06 | 2.59E+06 | 1.34E+06 |
| C04282 | 281.1403 | 17.458 | 66843 | 52567.68 | 5432.839 | 6253.807 | 5386.087 | 6031.568 | 68743.7 | 97255.2 |
| C04284 | 541.90516 | 21.846 | 161687 | 443404 | 2.89E+05 | 4.42E+05 | 3.19E+05 | 6.57E+05 | 365381 | 4.25E+05 |
| C04327 | 405.2401 | 10.616 | 1.40E+06 | 1.08E+06 | 7.02E+05 | 6.76E+05 | 8.76E+05 | 4.97E+05 | 12910.2 | 11820.8 |
| C04483 | 320.21993 | 14.688 | 141936 | 132871.8 | 14337.7 | 16363.06 | 18109.41 | 15244.95 | 16911.7 | 1.27E+05 |
| C04488 | 336.04166 | 18.693 | 27648.8 | 28977.72 | 5463.747 | 7071.687 | 5631.531 | 6656.264 | 73303.7 | 37253.2 |
| C04554 | 476.04515 | 10.464 | 16022.2 | 10181.89 | 3234.306 | 3554.541 | 3513.29 | 3209.917 | 39963.3 | 32288.5 |
| C04555 | 381.26373 | 17.339 | 107366 | 49780.98 | 37870.21 | 42948.87 | 53941.26 | 35656.99 | 116807 | 83998.3 |
| C04594 | 224.14047 | 12.321 | 23344.5 | 43193.33 | 6542.177 | 7814.08 | 11514.33 | 6493.254 | 108365 | 54909.4 |
| C04717 | 152.11977 | 12.57 | 103648 | 84497.37 | 10784.39 | 11292.14 | 11357.47 | 10849.78 | 277927 | 7.66E+05 |
| C04722 | 540.4973 | 16.028 | 55452.2 | 24084.48 | 26171.94 | 33574.44 | 18275 | 14212.24 | 125852 | 1.60E+05 |
| C04932 | 424.26605 | 10.658 | 1.96E+07 | 1.11E+07 | 5.31E+06 | 5.42E+06 | 4.49E+06 | 6.31E+06 | 110415 | 49791.3 |
| C05042 | 151.06301 | 3.062 | 101085 | 106137.7 | 92225.52 | 1.16E+05 | 1.62E+05 | 1.06E+05 | 3.13E+06 | 1.08E+07 |
| C05100 | 405.18391 | 10.585 | 14106.5 | 11538.57 | 2803.805 | 2935.692 | 13497.19 | 2760.407 | 32416.8 | 23312.1 |
| C05125 | 778.84024 | 1.634 | 15429.8 | 15557.54 | 17390.5 | 19354.87 | 17337.09 | 21421.66 | 36231.5 | 43157.8 |
| C05297 | 242.06758 | 14.51 | 90529.4 | 32524.3 | 9023.72 | 10069.27 | 9177.663 | 9458.51 | 58293.1 | 1.87E+05 |
| C05299 | 535.42729 | 14.629 | 432984 | 554872.1 | 4.04E+05 | 6.57E+05 | 3.72E+05 | 4.40E+05 | 30083.1 | 55756.4 |
| C05328 | 222.19765 | 15.03 | 23997.8 | 11163.82 | 10250.4 | 10701.35 | 12704.77 | 10245.9 | 98023.7 | 1.15E+05 |
| C05330 | 367.07958 | 12.087 | 4.77E+06 | 6.49E+06 | 3.58E+06 | 3.49E+06 | 3.10E+06 | 2.70E+06 | 127482 | 46953.2 |
| C05332 | 289.78302 | 17.715 | 79282.6 | 44266.64 | 4483.584 | 5218.135 | 4588.42 | 5443.662 | 212750 | 2.76E+05 |
| C05356 | 346.21104 | 10.541 | 47109.8 | 26051.5 | 10731.68 | 10681.38 | 11042.8 | 11946.39 | 23278.7 | 31207.3 |
| C05380 | 498.12459 | 17.966 | 168414 | 253847.9 | 29560.66 | 34408.87 | 28564.66 | 33444.66 | 1.53E+08 | 1.60E+08 |
| C05399 | 1108.6312 | 8.035 | 32373.1 | 20582.08 | 12127.46 | 17970.8 | 12227.38 | 5733.171 | 65770.6 | 36694.6 |
| C05455 | 464.31318 | 9.577 | 31343.1 | 14339.97 | 1729.06 | 1706.513 | 1936.674 | 1646.815 | 6316.23 | 6449.35 |
| C05465 | 200.02278 | 11.628 | 7.52E+07 | 1.47E+07 | 1.09E+07 | 1.05E+07 | 1.70E+07 | 1.08E+07 | 474189 | 1.99E+05 |
| C05466 | 563.2313 | 18.194 | 26180.1 | 26737.3 | 4519.728 | 5688.369 | 4450.687 | 5597.005 | 116940 | 95789.3 |
| C05475 | 985.65253 | 17.2 | 230831 | 110357.2 | 10143.03 | 11619.56 | 10011.07 | 11376.69 | 143190 | 1.55E+05 |
| C05498 | 336.66323 | 16.642 | 17504.9 | 21262.2 | 3074.54 | 3190.837 | 3453.175 | 2930.64 | 17976.6 | 11980 |
| C05516 | 336.21249 | 18.665 | 22004.2 | 25384.7 | 3997.127 | 5257.901 | 4022.827 | 4776.507 | 61822.2 | 1.00E+05 |
| C05543 | 1038.5775 | 12.65 | 40732.9 | 31756.37 | 3341.773 | 3790.725 | 3476.564 | 3513.162 | 80854.4 | 76364.5 |
| C05552 | 571.29106 | 22.151 | 53254.4 | 37976.31 | 6.51E+05 | 4.60E+05 | 1.26E+06 | 2.14E+05 | 70149.7 | 43459.8 |
| C05575 | 322.05381 | 12.126 | 72731.9 | 39486.02 | 22487.4 | 21814.77 | 69088.41 | 16400.71 | 191523 | 85149.4 |
| C05577 | 177.05487 | 8.714 | 47444.9 | 24747.36 | 5313.549 | 6141.627 | 5509.657 | 5875.312 | 404339 | 2.88E+05 |
| C05619 | 336.15891 | 1.944 | 1.62E+06 | 1.71E+06 | 1.29E+06 | 1.11E+06 | 1.18E+06 | 1.13E+06 | 9.93E+06 | 1.26E+07 |
| C05639 | 336.22344 | 11.356 | 47368.9 | 37730.69 | 6538.45 | 7055.592 | 6842.996 | 7156.852 | 331785 | 3.14E+05 |
| C05642 | 312.22766 | 12.699 | 56284.9 | 67891.28 | 13291.24 | 15882.06 | 14976.78 | 13981.9 | 321843 | 6.16E+05 |
| C05660 | 336.05351 | 10.625 | 72481.6 | 44755.64 | 12428.92 | 13262.2 | 13510.73 | 15049.13 | 65936.7 | 55402.1 |
| C05674 | 811.38871 | 9.534 | 15012.8 | 19022.2 | 2921.161 | 3107.348 | 3460.323 | 3005.109 | 19904.5 | 25267.3 |
| C05771 | 318.26151 | 13.372 | 2.24E+06 | 1.30E+06 | 1.31E+06 | 1.97E+06 | 1.22E+06 | 7.88E+05 | 48785.7 | 28948.3 |
| C05794 | 239.07608 | 1.482 | 994417 | 868765.7 | 4.19E+05 | 3.61E+05 | 3.51E+05 | 3.10E+05 | 236687 | 94826.8 |
| C05827 | 254.10503 | 10.588 | 139794 | 80646.69 | 22809.11 | 25235.94 | 25480.12 | 26988.63 | 180583 | 97748.3 |
| C05828 | 256.23962 | 10.911 | 148343 | 77517.06 | 18411.92 | 22535.93 | 19750.64 | 17787.66 | 89919.5 | 88240.5 |
| C05840 | 146.05832 | 17.871 | 5049.44 | 4374.251 | 4041.054 | 8095.752 | 3666.677 | 4309.355 | 19800.3 | 14184.7 |
| C05842 | 334.11753 | 14.198 | 36354.3 | 25740.35 | 6183.736 | 7536.597 | 8361.084 | 7269.86 | 122142 | 1.85E+05 |
| C05843 | 172.07205 | 1.499 | 26140.6 | 21344.25 | 3397.676 | 3430.892 | 3425.036 | 3367.091 | 25643.2 | 17928.9 |
| C05844 | 336.13185 | 1.882 | 163165 | 80173.11 | 20044.99 | 20575.02 | 19468.71 | 18787.17 | 136168 | 1.06E+05 |
| C05851 | 400.31759 | 16.491 | 343225 | 121425.3 | 8584.119 | 9821.581 | 9163.819 | 9832.264 | 42023.9 | 33282.1 |
| C05901 | 196.05871 | 3.1 | 44194.9 | 82408.62 | 60254.41 | 33157.26 | 1.51E+05 | 1.12E+05 | 68605 | 18199.5 |
| C05960 | 237.10773 | 12.168 | 67233.2 | 32477.43 | 3188.3 | 3547.649 | 3460.014 | 3310.085 | 29535.6 | 33303.9 |
| C05965 | 358.05037 | 11.252 | 57240.5 | 11256.84 | 3441.028 | 3744.196 | 3543.334 | 3540.051 | 4310.81 | 12456 |
| C05966 | 345.12062 | 8.531 | 12906.9 | 17866.09 | 12014.63 | 16252.5 | 4447.724 | 11857.39 | 42398.7 | 26669.6 |
| C06021 | 899.60735 | 12.368 | 168857 | 283447.5 | 35672.61 | 44285.09 | 39411.17 | 39730.94 | 1.82E+06 | 9.87E+05 |
| C06044 | 432.32106 | 7.865 | 164605 | 91891.1 | 21490.92 | 22384.54 | 1.22E+05 | 23781.27 | 150402 | 98919.3 |
| C06068 | 527.76791 | 2.158 | 1.04E+06 | 678159.1 | 5.38E+05 | 4.20E+05 | 5.12E+05 | 4.48E+05 | 282693 | 3.24E+05 |
| C06082 | 330.27604 | 13.326 | 61879.5 | 40624.77 | 9930.56 | 11132.56 | 11028.31 | 9058.52 | 1.34E+06 | 4.42E+05 |
| C06123 | 453.2825 | 13.835 | 22475.8 | 6430.539 | 2960.824 | 3077.206 | 4092.496 | 3137.156 | 17435.8 | 6024.28 |
| C06124 | 378.26242 | 11.912 | 1.4E+07 | 3267829 | 3311.88 | 3404.513 | 3562.928 | 3493.649 | 39915 | 17649.1 |
| C06178 | 569.36543 | 17.235 | 108490 | 69020.39 | 6309.54 | 7270.133 | 6402.383 | 7197.359 | 43417.9 | 35248.5 |
| C06213 | 569.36574 | 12.824 | 7720.22 | 7088.856 | 5408.713 | 8278.974 | 9005.954 | 5991.467 | 309411 | 52131.9 |
| C06244 | 216.17159 | 14.653 | 31817.7 | 29953.63 | 7189.474 | 8227.583 | 7042.532 | 8035.283 | 147998 | 1.93E+05 |
| C06323 | 281.90953 | 16.241 | 184533 | 106156.7 | 10412.91 | 11519.86 | 11113.6 | 11499.93 | 70798.6 | 82668 |
| C06345 | 326.14996 | 9.824 | 1.78E+06 | 742439.2 | 7.30E+05 | 6.83E+05 | 7.55E+05 | 5.51E+05 | 52353.5 | 47523.7 |
| C06390 | 239.22437 | 11.898 | 3.34E+07 | 1.09E+07 | 2.80E+06 | 2.98E+06 | 4.88E+06 | 2.74E+06 | 324942 | 25557.4 |
| C06414 | 326.28117 | 9.894 | 1.87E+06 | 806287.1 | 6.93E+05 | 6.18E+05 | 7.75E+05 | 6.15E+05 | 276788 | 55604.8 |
| C06424 | 234.16208 | 15.584 | 53544.4 | 45688.03 | 24249.27 | 22555.97 | 23082.58 | 20835.6 | 60658.3 | 66483.1 |
| C06429 | 582.31947 | 12.243 | 33550.9 | 22443.42 | 4768.226 | 5039.133 | 5403.074 | 4745.234 | 6135.19 | 16272.1 |
| C06517 | 452.24022 | 17.201 | 325520 | 166991.5 | 15018.58 | 17561.3 | 15114.45 | 16979.57 | 209970 | 2.27E+05 |
| C06528 | 280.22703 | 18.257 | 19465.3 | 24680.09 | 4333.418 | 5177.515 | 3934.464 | 5524.998 | 691193 | 6.39E+05 |
| C06567 | 365.25545 | 10.826 | 34391.5 | 22824.12 | 4655.305 | 5197.265 | 5140.974 | 5407.42 | 37819.5 | 1.14E+05 |
| C06575 | 215.97666 | 9.898 | 113400 | 97821.05 | 76152.85 | 1.40E+05 | 44328.54 | 57149.36 | 71271.7 | 61460.8 |
| C06635 | 729.69487 | 14.862 | 87444.6 | 41257.31 | 10702.72 | 11749.64 | 12999.43 | 11447.7 | 667844 | 1.59E+05 |
| C06654 | 337.20922 | 1.816 | 45449.4 | 53042.1 | 34114 | 46291.77 | 48329.57 | 88743.12 | 85854.5 | 79194.3 |
| C06747 | 300.17183 | 12.462 | 36370.8 | 49530.18 | 20739.41 | 43145.16 | 50247.71 | 30475.52 | 373943 | 36216.9 |
| C06804 | 558.32657 | 22.03 | 210580 | 158381.3 | 66073.93 | 1.07E+05 | 66010.25 | 1.12E+05 | 281001 | 2.88E+05 |
| C06894 | 562.28025 | 17.14 | 179994 | 82879.75 | 7971.848 | 9072.956 | 7850.232 | 8953.225 | 275176 | 3.69E+05 |
| C06905 | 281.161 | 18.077 | 27699.1 | 29092.8 | 3984.395 | 5050.323 | 4139.014 | 4941.933 | 79534.3 | 38673.9 |
| C06935 | 195.09699 | 6.435 | 245348 | 239964.5 | 1.61E+06 | 2.81E+05 | 3.42E+06 | 1.76E+06 | 126721 | 1.01E+05 |
| C06947 | 427.22171 | 10.056 | 241126 | 118789.8 | 25698.99 | 30057.42 | 27519.5 | 27588.78 | 288340 | 4.76E+05 |
| C06967 | 581.32106 | 11.96 | 88717 | 52646.26 | 4694.49 | 5161.895 | 5347.901 | 4692.168 | 27455.7 | 10520.8 |
| C06971 | 369.16685 | 10.909 | 473561 | 191443.2 | 34204.1 | 43917.69 | 37861.13 | 35555.4 | 1.76E+06 | 5.09E+05 |
| C06976 | 527.76795 | 2.238 | 256829 | 289899.3 | 1.91E+05 | 1.50E+05 | 1.77E+05 | 1.98E+05 | 1.72E+06 | 2.58E+06 |
| C06991 | 165.07148 | 2.745 | 3.99E+06 | 2.66E+06 | 1.24E+06 | 3.73E+06 | 1.22E+06 | 1.07E+06 | 463526 | 2.69E+05 |
| C06999 | 781.79235 | 3.563 | 4.26E+06 | 2.01E+06 | 6.40E+05 | 1.26E+06 | 4.14E+05 | 8.92E+05 | 60360.9 | 51382 |
| C07049 | 149.95577 | 1.846 | 48265.4 | 37964.12 | 46791.47 | 40599.94 | 37976.84 | 41372.89 | 110235 | 71999.9 |
| C07063 | 237.08364 | 2.827 | 45033.7 | 36586.88 | 18964.6 | 39104.07 | 57869.83 | 32642.06 | 126911 | 1.31E+05 |
| C07073 | 478.30744 | 11.634 | 34528 | 24619.55 | 3252.024 | 3419.457 | 3539.608 | 3234.595 | 6604.34 | 16722.7 |
| C07119 | 176.04703 | 13.583 | 265793 | 271381.1 | 34340.11 | 39332.06 | 36939.58 | 39050.22 | 5.09E+08 | 5.59E+08 |
| C07149 | 189.0421 | 13.307 | 158784 | 210871.9 | 25861.75 | 28787.96 | 28808.54 | 26967.95 | 6.59E+07 | 6.08E+07 |
| C07151 | 226.16774 | 9.481 | 154077 | 156862.9 | 22035.07 | 23656.76 | 24714.43 | 23224.51 | 1.27E+07 | 1.11E+07 |
| C07202 | 330.23974 | 11.26 | 37182.8 | 25491.41 | 5015.613 | 5344.916 | 5616.998 | 5370.761 | 185624 | 2.48E+05 |
| C07203 | 482.31941 | 11.705 | 59169.4 | 23172.81 | 3868.449 | 4603.247 | 3994.956 | 3284.772 | 18050.1 | 48321.7 |
| C07219 | 326.28131 | 19.337 | 69471.1 | 34183.44 | 3717.153 | 3813.006 | 3862.52 | 3739.281 | 8698.82 | 13349.3 |
| C07251 | 519.33197 | 17.137 | 491364 | 226983.8 | 21818.23 | 24928.78 | 21496.64 | 24448.81 | 6.02E+07 | 7.02E+07 |
| C07264 | 333.06431 | 19.975 | 112574 | 81363.26 | 20391 | 25036.17 | 22423.68 | 25753.56 | 2.06E+06 | 1.19E+06 |
| C07272 | 176.04701 | 11.537 | 171086 | 161940.4 | 28013.98 | 31067.46 | 32706.53 | 32630.75 | 6.67E+07 | 6.16E+07 |
| C07276 | 495.3311 | 11.756 | 283691 | 89609.96 | 3.31E+05 | 3.94E+05 | 4.57E+05 | 1.42E+05 | 431134 | 1.52E+05 |
| C07289 | 330.23987 | 13.072 | 215572 | 109423.1 | 22962.38 | 24750.67 | 24798.03 | 22988.89 | 624960 | 7.27E+05 |
| C07325 | 354.13716 | 11.289 | 51969.2 | 34465.46 | 6187.662 | 6714.146 | 6681.345 | 6793.817 | 3.14E+06 | 2.50E+06 |
| C07327 | 99.06871 | 2.891 | 443546 | 280052.3 | 38349.62 | 42623.22 | 40359.81 | 44078.94 | 5.06E+06 | 4.61E+06 |
| C07370 | 164.04702 | 2.226 | 79753.2 | 68808.49 | 17065.55 | 17861.67 | 18787.59 | 17915.98 | 2.48E+06 | 2.86E+06 |
| C07440 | 362.20867 | 11.672 | 46544.8 | 21515.75 | 3332.827 | 3549.697 | 3728.263 | 3220.464 | 11739.6 | 28149.2 |
| C07481 | 164.09854 | 20.179 | 65022.6 | 53791.57 | 22768.73 | 48708.33 | 20791.81 | 36558.58 | 74483.6 | 57297.9 |
| C07493 | 237.8547 | 1.926 | 1.27E+06 | 668429.2 | 4.11E+05 | 3.77E+05 | 3.79E+05 | 4.67E+05 | 100730 | 1.23E+05 |
| C07496 | 559.29346 | 17.602 | 57115.2 | 61303.81 | 5733.234 | 6862.642 | 5836.866 | 6705.489 | 460902 | 5.41E+05 |
| C07527 | 278.14877 | 2.453 | 11646.7 | 9757.089 | 15020.47 | 11450.92 | 11162.21 | 18123.64 | 65523.4 | 59057.7 |
| C07577 | 430.30711 | 7.684 | 193378 | 82167.41 | 90057.82 | 79080.26 | 85016.49 | 89574.09 | 89337.7 | 42559.9 |
| C07585 | 252.07194 | 6.734 | 26123.1 | 26543.2 | 8778.484 | 6663.542 | 6559.731 | 7024.525 | 182917 | 30711.5 |
| C07592 | 163.05142 | 5.073 | 1.05E+07 | 3.92E+06 | 3.22E+06 | 3.58E+06 | 2.98E+06 | 3.26E+06 | 512695 | 3.19E+05 |
| C07610 | 95.03669 | 12.997 | 20818.4 | 32384.09 | 2615.047 | 2979.017 | 3291.749 | 2824.324 | 360035 | 3.77E+05 |
| C07652 | 235.97694 | 7.775 | 164629 | 159500.3 | 1.17E+05 | 5.37E+05 | 9.11E+05 | 4.93E+05 | 208588 | 1.45E+05 |
| C07669 | 346.21123 | 15.048 | 28587.1 | 24332.83 | 6446.362 | 9872.007 | 23076.39 | 6798.663 | 41976.8 | 21480.3 |
| C07712 | 457.33373 | 9.561 | 39063.4 | 17989.19 | 6821.561 | 12838.03 | 14437.09 | 10936.02 | 198829 | 62238.4 |
| C07713 | 498.99138 | 18.505 | 54230 | 58626.05 | 11314.75 | 13991.29 | 11193.5 | 12847.33 | 1.55E+07 | 1.17E+07 |
| C07818 | 197.0103 | 3.598 | 54283.6 | 39743.7 | 11722.73 | 10224.71 | 11579.06 | 12191.09 | 52557.1 | 1.71E+05 |
| C07844 | 227.66438 | 16.542 | 690227 | 240752 | 18171.03 | 20358.56 | 19400.21 | 20651.52 | 325862 | 2.13E+05 |
| C07868 | 330.21822 | 10.758 | 65475.3 | 27458.64 | 11142.76 | 12715.88 | 13465.03 | 15429.92 | 338831 | 4.04E+05 |
| C07895 | 558.32661 | 11.11 | 64412.9 | 37542.09 | 6670.168 | 6971.973 | 7046.3 | 7147 | 172552 | 1.20E+05 |
| C08169 | 176.0472 | 1.137 | 122165 | 98376.95 | 16318.68 | 18179.82 | 16145.16 | 17655.07 | 4.13E+07 | 2.26E+07 |
| C08261 | 519.33164 | 15.617 | 105149 | 159061.9 | 60866.82 | 72392.28 | 69197.52 | 54286.72 | 960002 | 1.78E+05 |
| C08287 | 151.89272 | 1.941 | 2.58E+06 | 2.72E+06 | 1.64E+06 | 2.08E+06 | 1.77E+06 | 1.75E+06 | 968226 | 7.29E+05 |
| C08288 | 495.33109 | 11.758 | 209573 | 94296.04 | 3.05E+05 | 2.01E+05 | 2.68E+05 | 1.21E+05 | 128847 | 42713.1 |
| C08302 | 336.04197 | 2.022 | 162762 | 90763.29 | 62403.06 | 62699.63 | 60762.25 | 65329.55 | 122237 | 1.91E+05 |
| C08316 | 278.07835 | 17.237 | 34019.4 | 27470.97 | 3115.479 | 3128.929 | 3400.031 | 2966.276 | 6238.43 | 6835.53 |
| C08317 | 237.82321 | 1.982 | 43475.9 | 74498.07 | 15253.68 | 15053.56 | 14217.07 | 12032.89 | 220793 | 2.04E+05 |
| C08362 | 140.04715 | 1.354 | 45518.5 | 36391.16 | 4816.258 | 5373.204 | 4792.029 | 5681.079 | 62161.4 | 44079.2 |
| C08363 | 186.12437 | 1.192 | 288899 | 229079.4 | 37758.65 | 42785.67 | 42108.48 | 41249.85 | 954785 | 1.98E+06 |
| C08365 | 132.08966 | 2.711 | 802955 | 371400 | 63624.63 | 65107.13 | 65273.71 | 73364.75 | 471047 | 3.82E+05 |
| C08367 | 265.11568 | 13.716 | 54919.9 | 57909.21 | 7401.512 | 8470.751 | 7953.672 | 8503.016 | 2.82E+06 | 3.44E+06 |
| C08397. | 336.22694 | 1.752 | 4.68E+06 | 5.02E+06 | 3.47E+06 | 3.49E+06 | 2.94E+06 | 4.20E+06 | 436125 | 4.18E+05 |
| C08441 | 193.11413 | 9.432 | 88517.6 | 85436.55 | 12537.07 | 13204.42 | 13605.98 | 12671.54 | 684359 | 6.60E+05 |
| C08497 | 266.1877 | 16.557 | 44849.2 | 30434.63 | 3587.977 | 3689.782 | 4285.471 | 3433.91 | 30798.2 | 33014 |
| C08588 | 234.03554 | 16.771 | 193568 | 106650.2 | 9292.066 | 10665.93 | 9745.013 | 10303.72 | 256375 | 1.49E+05 |
| C08859 | 196.05835 | 7.346 | 30148.2 | 24477.34 | 8010.158 | 8409.834 | 9614.263 | 9378.009 | 924491 | 8.21E+05 |
| C08876 | 527.76334 | 19.272 | 71133.2 | 57981.65 | 19237.72 | 20898.76 | 15887.49 | 19929.02 | 6.18E+06 | 3.61E+06 |
| C08994 | 337.33318 | 10.773 | 67569.7 | 61815.88 | 22916.66 | 65406.85 | 91382.53 | 35892.72 | 170202 | 58506 |
| C09069 | 561.30844 | 18.05 | 50858.9 | 45215.85 | 6563.274 | 8790.966 | 6472.777 | 7940.286 | 925100 | 1.22E+06 |
| C09210 | 931.43677 | 12.374 | 53741.6 | 112169.9 | 11220.54 | 13506.75 | 13169.95 | 12340.59 | 346799 | 3.94E+05 |
| C09226 | 234.16215 | 7.592 | 69201.7 | 138364.8 | 1.08E+05 | 2.80E+05 | 4.49E+05 | 2.92E+05 | 135123 | 1.28E+05 |
| C09311 | 95.03738 | 12.656 | 50572.7 | 41638.19 | 4106.791 | 4710.226 | 4248.999 | 4455.769 | 238232 | 2.41E+05 |
| C09315 | 318.26153 | 14.467 | 1.12E+06 | 596418.2 | 2.32E+05 | 3.43E+05 | 6.99E+05 | 3.04E+05 | 308267 | 3.34E+05 |
| C09622 | 570.3719 | 13.641 | 25448.9 | 39993.54 | 59728.19 | 55084.89 | 56406.63 | 4.58E+05 | 30830.2 | 27649.3 |
| C09662 | 428.14316 | 11.615 | 40008.6 | 46382.86 | 7567.64 | 8294.512 | 9267.947 | 8406.211 | 577571 | 69667.3 |
| C09715 | 132.07743 | 6.765 | 288833 | 310748.4 | 1.28E+05 | 87438.49 | 97390.43 | 92829.19 | 668942 | 1.14E+06 |
| C09861 | 280.99627 | 22.094 | 243878 | 82231.46 | 3.82E+05 | 2.91E+05 | 4.38E+05 | 3.48E+05 | 245001 | 1.17E+05 |
| C09947 | 237.9979 | 1.973 | 468476 | 271742.6 | 1.00E+05 | 93638.29 | 75868.06 | 89987.9 | 117579 | 2.15E+05 |
| C09952 | 254.10492 | 10.293 | 15890.4 | 11599.51 | 6607.247 | 7426.315 | 6208.41 | 8103.695 | 27615.3 | 27549.7 |
| C10030 | 541.33204 | 14.598 | 1.65E+07 | 2.27E+07 | 6.46E+06 | 1.13E+07 | 7.13E+06 | 8.41E+06 | 14634.5 | 29055.8 |
| C10056 | 778.84703 | 17.395 | 118078 | 65977.78 | 6259.009 | 7517.186 | 6479.295 | 7250.599 | 29442.1 | 40433.6 |
| C10138 | 281.13999 | 10.491 | 175399 | 118043.5 | 16004.41 | 17026.72 | 16652.99 | 15198.31 | 273206 | 3.97E+05 |
| C10199 | 311.01088 | 12.926 | 327060 | 300154.5 | 2.16E+05 | 2.76E+05 | 1.52E+05 | 1.23E+05 | 17597 | 27235.9 |
| C10387 | 164.04701 | 2.351 | 98727.7 | 98448.32 | 18990.64 | 21490.12 | 18272.91 | 21571.71 | 4.10E+06 | 2.05E+06 |
| C10457 | 151.99305 | 1.862 | 327131 | 496593.7 | 1.37E+05 | 99506.76 | 1.11E+05 | 1.13E+05 | 173842 | 1.28E+05 |
| C10462 | 330.23991 | 12.733 | 32313.7 | 31124.11 | 10270.91 | 12865.84 | 13727.7 | 10534.33 | 1.24E+06 | 2.04E+06 |
| C10858 | 531.30333 | 2.011 | 572459 | 383050.8 | 99646.53 | 1.34E+05 | 1.47E+05 | 1.26E+05 | 3.07E+07 | 1.78E+07 |
| C10947 | 226.16782 | 9.707 | 539369 | 317718.3 | 1.04E+05 | 92953.92 | 1.00E+05 | 90451.76 | 5.25E+06 | 2.97E+06 |
| C10996 | 220.09404 | 6.983 | 297080 | 336576.4 | 68297.23 | 67865.45 | 66210.86 | 72321.52 | 898910 | 4.23E+05 |
| C10997 | 98.52408 | 1.461 | 54268.6 | 37484.41 | 24149.93 | 17283.72 | 25533.31 | 24683.35 | 5.35E+07 | 4.09E+07 |
| C11002 | 101.08446 | 1.828 | 191106 | 66809.21 | 12704.65 | 13837.02 | 12826.7 | 13021.88 | 562707 | 6.06E+05 |
| C11005 | 84.02018 | 1.595 | 1.92E+06 | 914271.8 | 4.11E+05 | 5.13E+05 | 4.11E+05 | 4.29E+05 | 144564 | 2.44E+05 |
| C11011 | 345.28526 | 11.45 | 33906.2 | 24084.14 | 3685.664 | 4015.314 | 3738.464 | 4225.735 | 92821.6 | 1.83E+05 |
| C11077 | 157.90158 | 1.861 | 127834 | 89935.07 | 83885.28 | 69179.96 | 74215.09 | 73928.18 | 174951 | 1.09E+05 |
| C11087 | 166.02641 | 12.8 | 236742 | 234551.9 | 22550.58 | 25262.71 | 27859.5 | 24571.48 | 3.62E+09 | 5.32E+09 |
| C11135 | 239.15351 | 9.712 | 188937 | 267087.9 | 2.30E+05 | 3.81E+05 | 1.34E+05 | 1.95E+05 | 2.40E+06 | 2.31E+06 |
| C11156 | 145.01375 | 1.803 | 99070.5 | 43550.82 | 8093.022 | 8740.956 | 8093.089 | 8459.246 | 171792 | 68213.6 |
| C11185 | 129.04241 | 1.876 | 679972 | 436102.2 | 95812.19 | 93364.27 | 86537.98 | 90090.47 | 1.76E+06 | 1.59E+06 |
| C11284 | 328.16558 | 9.847 | 4.74E+07 | 1.08E+07 | 5.64E+06 | 4.88E+06 | 5.24E+06 | 4.98E+06 | 36468.5 | 30188.1 |
| C11409 | 150.10422 | 8.921 | 64523.9 | 77684.64 | 7902.069 | 8469.344 | 9410.009 | 8289.891 | 163178 | 2.12E+05 |
| C11434 | 552.14897 | 22.13 | 641331 | 1.13E+06 | 3.38E+05 | 4.34E+05 | 1.10E+05 | 6.20E+05 | 469373 | 2.76E+05 |
| C11458 | 227.66459 | 16.208 | 280846 | 192227.6 | 18693.24 | 21511.36 | 19867 | 20773.37 | 4.45E+06 | 5.20E+06 |
| C11527 | 247.21468 | 8.102 | 38722 | 19766.63 | 6139.169 | 6284.359 | 6599.153 | 5950.151 | 263059 | 1.64E+05 |
| C11695 | 335.19414 | 10.562 | 45614.4 | 11778.71 | 20689.94 | 18745.04 | 26522.33 | 14689.86 | 216991 | 31530.6 |
| C11787 | 519.33176 | 16.991 | 319074 | 240879.3 | 23135.16 | 25697.65 | 24280.79 | 25925.1 | 2.49E+07 | 2.52E+07 |
| C11837 | 525.75272 | 15.147 | 180293 | 131767.3 | 24103.98 | 28361.35 | 26579.56 | 28245.94 | 464554 | 2.02E+05 |
| C11853 | 346.23463 | 18.853 | 94331.4 | 57150.08 | 5119.751 | 5546.371 | 6340.869 | 5255.057 | 11962.3 | 15137.1 |
| C11857 | 348.17529 | 13.552 | 28811.4 | 30445.48 | 4312.435 | 4367.389 | 4813.388 | 4103.14 | 15938.1 | 27203.4 |
| C11859 | 346.21354 | 17.936 | 54824.3 | 44110.23 | 4653.586 | 4631.835 | 4898.58 | 4471.356 | 50272.7 | 47310 |
| C11918 | 176.0469 | 12.813 | 310828 | 266175.5 | 30225.57 | 33287.52 | 36977.48 | 32378.23 | 3.91E+09 | 5.62E+09 |
| C12026 | 149.01898 | 12.812 | 296553 | 353055.8 | 1.48E+05 | 1.60E+05 | 1.33E+05 | 1.30E+05 | 3.73E+08 | 4.98E+08 |
| C12110 | 282.92238 | 21.955 | 44620.2 | 25421.88 | 1.00E+05 | 59889.97 | 1.30E+05 | 58800.54 | 283225 | 3.61E+05 |
| C12144 | 196.05848 | 2.879 | 530997 | 608917.7 | 2.69E+05 | 2.32E+05 | 1.88E+05 | 2.57E+05 | 36619.5 | 46922.8 |
| C12272 | 495.33171 | 15.9 | 401274 | 406838.4 | 40980.65 | 44493.77 | 43924.18 | 44842.83 | 780971 | 3.27E+05 |
| C12448 | 530.16119 | 2.171 | 1.55E+07 | 2.13E+07 | 1.57E+07 | 1.41E+07 | 1.88E+07 | 1.43E+07 | 437994 | 5.09E+05 |
| C12456 | 251.10358 | 7.701 | 102408 | 78378.68 | 14649.87 | 14041.97 | 13556.18 | 17507.99 | 277832 | 97834.8 |
| C12476 | 155.90469 | 2.097 | 126198 | 150287.1 | 68951.18 | 58772.75 | 60804.72 | 86155.18 | 75409.3 | 56047.9 |
| C12535 | 190.09496 | 13.641 | 236146 | 263349.5 | 29192.25 | 33768.7 | 30928.59 | 32703.76 | 1.58E+07 | 3.23E+07 |
| C12634 | 166.09898 | 10.241 | 137836 | 67854.21 | 16871.04 | 17977.56 | 18204.23 | 17144.12 | 468806 | 1.91E+05 |
| C12849 | 337.04283 | 14.108 | 21930.9 | 21184.3 | 3268.755 | 4225.985 | 4009.785 | 3794.115 | 118260 | 55410.3 |
| C13179 | 782.19597 | 2.809 | 82077.2 | 39174.15 | 16558.31 | 19545.42 | 19735.11 | 18230.69 | 5.27E+06 | 2.94E+06 |
| C13453 | 328.15135 | 17.342 | 88669.9 | 63711.05 | 7693.63 | 7478.93 | 8088.611 | 7187.377 | 12931.2 | 1.38E+05 |
| C13690 | 374.9453 | 10.665 | 179115 | 96901.02 | 82821.42 | 3.89E+05 | 1.96E+05 | 1.53E+05 | 129497 | 93981.7 |
| C13747 | 193.07371 | 1.73 | 139316 | 84296.09 | 18243.08 | 18324.39 | 17260.59 | 17439.2 | 63280.5 | 96292 |
| C14150 | 454.10425 | 14.417 | 62823.7 | 111478.5 | 40810.42 | 65902.11 | 78229.64 | 71435.47 | 24908 | 15775.6 |
| C14155 | 280.925 | 22.302 | 190441 | 196392.4 | 5.24E+05 | 3.22E+05 | 4.70E+05 | 3.19E+05 | 201500 | 2.00E+05 |
| C14175 | 156.91739 | 1.851 | 143644 | 94368.76 | 24478.63 | 26276.4 | 26116.76 | 25594.8 | 243672 | 1.74E+05 |
| C14211 | 729.69896 | 5.069 | 4763.19 | 5191.983 | 6027.314 | 6380.956 | 6179.069 | 6594.184 | 86420.3 | 58132.4 |
| C14214 | 519.33114 | 14.881 | 97202.8 | 167511.2 | 43839.63 | 74346.11 | 39791.84 | 29565.76 | 35635.7 | 25683.2 |
| C14279 | 96.0841 | 12.611 | 115402 | 89992.02 | 11161.09 | 12405.29 | 11654.73 | 11457.37 | 336154 | 4.74E+05 |
| C14291 | 559.31073 | 18.953 | 34494.8 | 32889.22 | 6243.123 | 7615.115 | 6068.428 | 7561.123 | 125575 | 74037.5 |
| C14302 | 507.32509 | 10.407 | 48593.8 | 24800.17 | 2916.794 | 3165.037 | 3134.497 | 2832.559 | 39388.1 | 85536.6 |
| C14323 | 569.36557 | 11.532 | 87623.1 | 45615.17 | 1.74E+05 | 2.20E+05 | 1.36E+05 | 81721.5 | 56277.6 | 14033.9 |
| C14331 | 415.32543 | 9.542 | 19723.9 | 4002.278 | 7736.167 | 41381.4 | 70835.62 | 27689.83 | 35301.4 | 15528.5 |
| C14337 | 338.24528 | 20.232 | 66404.2 | 69711.43 | 16484.19 | 18338.73 | 16105.45 | 18784.96 | 625682 | 9.17E+05 |
| C14387 | 364.07878 | 10.066 | 31194.4 | 15335.75 | 3648.72 | 3877.66 | 3882.804 | 3906.363 | 112494 | 46820 |
| C14439 | 298.24997 | 16.261 | 135779 | 188746.9 | 43176.69 | 47155.97 | 55386.98 | 42906.87 | 603498 | 2.35E+07 |
| C14469 | 509.34704 | 11.236 | 216940 | 108330.6 | 2.86E+05 | 4.27E+05 | 1.56E+05 | 96107.98 | 37884.7 | 2.59E+06 |
| C14516 | 237.88876 | 1.637 | 169196 | 85965.51 | 20297.25 | 21284.03 | 20623.09 | 20091.92 | 421034 | 4.98E+05 |
| C14602 | 247.1452 | 6.729 | 183194 | 101709.4 | 59654.74 | 86449.61 | 39087.02 | 59743.86 | 351974 | 61244.8 |
| C14686 | 569.36833 | 15.973 | 320644 | 386751.2 | 93826.99 | 1.46E+05 | 99982.82 | 1.07E+05 | 11836.9 | 25480.8 |
| C14695 | 496.98186 | 17.602 | 136189 | 215367.9 | 25621.53 | 28572.73 | 23228.3 | 26959.32 | 1.41E+07 | 1.50E+07 |
| C14772 | 447.29567 | 12.548 | 8587.21 | 6156.931 | 6846.428 | 11512.7 | 14460.42 | 9929.066 | 28835.1 | 20679.9 |
| C14773 | 450.05082 | 11.884 | 55636.8 | 35052.2 | 2922.482 | 3297.672 | 3412.889 | 2949.587 | 4889.39 | 8179.05 |
| C14774 | 479.29683 | 14.659 | 56158.8 | 43286.39 | 4650.517 | 5214.726 | 5782.123 | 4834.361 | 14180.4 | 11982.7 |
| C14775 | 479.29694 | 9.783 | 167867 | 55752.04 | 3803.961 | 4350.888 | 3597.415 | 4221.796 | 7637.01 | 5567.13 |
| C14781 | 344.13229 | 14.651 | 16777.7 | 6721.992 | 4421.014 | 4538.827 | 5091.653 | 3786.355 | 6720.92 | 5464.08 |
| C14812 | 354.24352 | 9.954 | 15470.5 | 12201.91 | 3558.712 | 9547.916 | 3774.976 | 5763.316 | 15894.7 | 13824 |
| C14813 | 361.10732 | 1.382 | 43362.5 | 83598.94 | 18525.57 | 27882.95 | 22183.84 | 29274.75 | 71396.7 | 20984 |
| C14823 | 421.13109 | 11.556 | 36175.3 | 25129.05 | 4375.936 | 4790.896 | 4919.829 | 4389.936 | 8683.33 | 11374.4 |
| C14825 | 152.1197 | 14.126 | 387277 | 252113.9 | 58022.88 | 74996.01 | 74629.24 | 66066.72 | 1.05E+06 | 1.07E+06 |
| C14832 | 330.23986 | 14.031 | 32362.7 | 35242.7 | 6110.37 | 6590.532 | 6386.511 | 6165.392 | 576224 | 2.11E+05 |
| C14833 | 142.09906 | 4.834 | 242824 | 226235.7 | 56600.26 | 67274.29 | 66127.75 | 1.37E+05 | 7.50E+06 | 6.63E+05 |
| C14871 | 307.87065 | 1.666 | 207908 | 93588.43 | 5066.977 | 5321.702 | 6005.623 | 6945.341 | 5562843 | 1398625 |
| C15025 | 330.23977 | 13.653 | 467187 | 255330.8 | 35441.55 | 39202.24 | 38774.03 | 37554.67 | 8.72E+06 | 9.11E+06 |
| C15501 | 247.14571 | 7.399 | 24649 | 18854.52 | 8308.495 | 8023.849 | 9403.26 | 8813.846 | 502059 | 3.74E+05 |
| C15502 | 247.21427 | 6.875 | 22548.5 | 16009.53 | 7639.499 | 18473.26 | 8105.078 | 7108.437 | 347673 | 1.77E+05 |
| C15542 | 581.32061 | 12.116 | 207211 | 87311.05 | 8646.911 | 9140.476 | 8925.931 | 8500.289 | 31184.2 | 49275.3 |
| C15607 | 756.46126 | 9.887 | 24989.9 | 21518.72 | 5401.57 | 4753.214 | 5425.636 | 4682.889 | 6946.97 | 17412.8 |
| C15986 | 674.23843 | 14.213 | 48351 | 91328.47 | 1.01E+05 | 3.78E+05 | 4.57E+05 | 2.46E+05 | 29687.3 | 44076.3 |
| C15989 | 224.14039 | 11.894 | 53180.2 | 39597.53 | 21698.08 | 16957.61 | 14340.72 | 20767.4 | 353495 | 4.50E+05 |
| C16147 | 198.02416 | 1.485 | 7.13E+06 | 6.78E+06 | 1.89E+06 | 2.25E+06 | 2.05E+06 | 1.95E+06 | 780744 | 4.33E+05 |
| C16311 | 146.10526 | 1.505 | 224878 | 90357.01 | 24450.39 | 25096.26 | 26250.31 | 25344.6 | 2.49E+06 | 1.79E+06 |
| C16322 | 335.82888 | 17.921 | 31709 | 34606.18 | 4408.309 | 5424.941 | 4439.175 | 5386.507 | 40465.5 | 64551.3 |
| C16323 | 148.06605 | 3.792 | 73635.6 | 113950.2 | 72256.05 | 5.45E+05 | 7.10E+05 | 2.19E+05 | 266129 | 1.48E+05 |
| C16360 | 238.94815 | 10.701 | 5308.43 | 6495.508 | 1.32E+05 | 1.15E+05 | 10071.68 | 12925.98 | 90474 | 40316.9 |
| C16503 | 189.04209 | 11.094 | 717636 | 471407.9 | 2.55E+05 | 3.46E+05 | 2.56E+05 | 1.76E+05 | 25950.6 | 26171.8 |
| C16512 | 330.2398 | 12.993 | 59514.8 | 67737.14 | 15842.7 | 20017.45 | 17015.79 | 16564.79 | 2.40E+06 | 3.24E+06 |
| C16513 | 266.18778 | 14.702 | 41814.4 | 29254 | 3249.48 | 3545.989 | 4090.975 | 3367.105 | 16100 | 3975.71 |
| C16522 | 158.05748 | 20.098 | 29982.5 | 24257.33 | 5465.181 | 6624.568 | 5354.697 | 6444.855 | 28221.9 | 45870 |
| C16526 | 158.05765 | 7.757 | 763795 | 528112.6 | 1.26E+05 | 1.22E+05 | 1.18E+05 | 1.35E+05 | 3.52E+06 | 2.45E+06 |
| C16538 | 238.0883 | 10.571 | 5327.92 | 6490.556 | 1.32E+05 | 1.15E+05 | 1.39E+05 | 93029.68 | 243425 | 33982.3 |
| C16577 | 935.55559 | 15.526 | 29149.9 | 32818.01 | 7701.876 | 8182.836 | 13601.12 | 7882.336 | 68144.7 | 88550.2 |
| C16582 | 138.04391 | 1.45 | 15007.1 | 7641.164 | 5933.873 | 12027.45 | 5811.891 | 10157.14 | 220478 | 1.68E+05 |
| C16584 | 133.06312 | 0.886 | 2370.36 | 1768.144 | 1898.184 | 1979.513 | 1995.11 | 1983.84 | 36307.3 | 18446.8 |
| C16589 | 454.32646 | 14.437 | 118378 | 106784.7 | 12942.92 | 14143.04 | 15938.29 | 13433.38 | 32027.6 | 48375.3 |
| C16607 | 674.23844 | 1.639 | 61211.6 | 35993.34 | 17451.97 | 22899.47 | 20046.3 | 22121.53 | 41059.2 | 32689.2 |
| C16614 | 82.00831 | 1.909 | 54236.5 | 31037.71 | 8502.139 | 8089.444 | 8180.241 | 7269.126 | 59305.7 | 58230.9 |
| C16634 | 242.192 | 11.663 | 39127 | 24888.9 | 2990.824 | 3160.461 | 3489.475 | 3009.112 | 12887.2 | 15954.9 |
| C16635 | 320.06389 | 10.76 | 16304.1 | 11921.8 | 6641.303 | 5660.543 | 2854.053 | 3694.207 | 26383 | 17225 |
| C16660 | 476.2033 | 2.301 | 39329.7 | 24209.55 | 35117.19 | 41008.73 | 44969.78 | 54383.96 | 187602 | 1.34E+05 |
| C16665 | 377.18129 | 2.003 | 1.99E+06 | 1.92E+06 | 6.28E+05 | 8.21E+05 | 5.33E+05 | 9.57E+05 | 138522 | 2.36E+05 |
| C16712 | 237.03789 | 2.158 | 378418 | 148767.2 | 50393.8 | 45926.5 | 47252.72 | 46311.34 | 201286 | 1.92E+05 |
| C16715 | 482.35667 | 14.602 | 1.54E+07 | 2.80E+07 | 6.65E+06 | 1.17E+07 | 7.39E+06 | 8.66E+06 | 29099.1 | 58613.8 |
| C16759 | 337.33373 | 10.549 | 39379.7 | 26267.62 | 13730 | 35528.02 | 35445.19 | 19485.79 | 220450 | 31073.6 |
| C16765 | 362.20854 | 9.587 | 56592.6 | 35350.93 | 3.70E+05 | 4.00E+05 | 16893.95 | 20489.81 | 39863 | 23959.8 |
| C16930 | 562.28054 | 20.16 | 27839.4 | 27970.83 | 6109.691 | 6818.2 | 5816.045 | 6906.145 | 264059 | 1.89E+05 |
| C17267 | 400.27937 | 14.931 | 10093.7 | 14562.02 | 10256.85 | 18887.01 | 13771.25 | 9705.229 | 12385 | 10305.2 |
| C17331 | 495.33099 | 9.172 | 2930.42 | 2566.421 | 2224.463 | 2103.253 | 1849.187 | 3855.714 | 2439.43 | 9303.22 |
| C17332 | 487.23726 | 15.129 | 24404 | 14631.34 | 11162.57 | 15927.08 | 9850.378 | 7064.553 | 34317.6 | 18825.3 |
| C17333 | 476.20381 | 11.671 | 21670.1 | 20893.82 | 4307.273 | 3862.073 | 7132.22 | 3645.276 | 37471.7 | 15027 |
| C17336 | 478.21878 | 12.763 | 6691.71 | 8322.659 | 4225.699 | 5841.354 | 6440.474 | 4515.414 | 13741.8 | 11375.1 |
| C17339 | 462.14184 | 9.519 | 2370.59 | 2549.683 | 3081.226 | 9185.212 | 15552.38 | 8153.745 | 9654.45 | 8305.69 |
| C17367 | 281.64244 | 17.222 | 101065 | 57946.65 | 5704.242 | 6608.546 | 5799.895 | 6547.778 | 42821.4 | 29850.8 |
| C17590 | 561.30878 | 18.46 | 14556.6 | 23347.03 | 4641.756 | 5645.022 | 4402.249 | 5530.628 | 155501 | 1.12E+05 |
| C17751 | 462.22244 | 19.792 | 26927.5 | 21813.25 | 5571.827 | 7062.925 | 5699.553 | 6917.155 | 37733.9 | 67283.3 |
| C18034 | 531.3036 | 2.235 | 734569 | 589703.3 | 1.39E+05 | 1.59E+05 | 1.76E+05 | 1.31E+05 | 9.19E+07 | 8.53E+07 |
| C18043 | 778.84017 | 1.646 | 21595 | 17957.18 | 21765.78 | 22545.9 | 20186.93 | 25835.88 | 104796 | 66557.3 |
| C18044 | 442.00819 | 16.246 | 32664.8 | 22419.42 | 4046.303 | 3858.07 | 4929.961 | 3633.86 | 6392.88 | 13736.5 |
| C18049 | 146.10515 | 2.643 | 591898 | 301963.1 | 45968.63 | 46686.25 | 48258.45 | 49416.64 | 1.92E+06 | 1.79E+06 |
| C18403 | 164.04704 | 3.498 | 654679 | 707615.6 | 1.39E+05 | 1.44E+05 | 1.35E+05 | 1.45E+05 | 7.75E+06 | 5.08E+06 |
| C18417 | 410.22784 | 11.054 | 164824 | 305294.5 | 91166.85 | 97765.16 | 65738.33 | 1.12E+05 | 280013 | 1.25E+05 |
| C18421 | 224.14033 | 17.515 | 76665 | 71412.48 | 6795.691 | 7950.014 | 6733.007 | 7750.142 | 65626.7 | 47549 |
| C18432 | 239.14056 | 3.767 | 253726 | 194866 | 82671.49 | 1.17E+05 | 88484.31 | 84621.91 | 266437 | 1.92E+05 |
| C18433 | 238.08336 | 1.899 | 110012 | 58358.02 | 21178.09 | 19212.85 | 17312.31 | 18629.38 | 110456 | 79397.2 |
| C18786 | 374.22512 | 10.803 | 135292 | 130396.9 | 1.65E+05 | 4.26E+05 | 3.35E+05 | 2.48E+05 | 219634 | 1.52E+05 |
| C18796 | 541.33229 | 22.37 | 188447 | 288216.7 | 1.45E+05 | 1.78E+05 | 2.16E+05 | 1.92E+05 | 288539 | 3.21E+05 |
| C18868 | 521.34751 | 17.968 | 122791 | 126368.2 | 15859.81 | 19560.14 | 16095.13 | 18806.59 | 1.07E+07 | 3.93E+06 |
| C18912 | 283.65808 | 22.302 | 78611.5 | 62117.77 | 2.35E+05 | 1.21E+05 | 2.01E+05 | 1.24E+05 | 141755 | 1.60E+05 |
| C18980 | 507.32835 | 11.33 | 12453.8 | 19368.85 | 3639.654 | 3879.706 | 3848.53 | 3769.069 | 12368.3 | 16249.8 |
| C19015 | 132.07775 | 6.349 | 716458 | 346303 | 1.80E+05 | 2.05E+05 | 1.10E+05 | 85376.33 | 305206 | 3.93E+05 |
| C19035 | 482.2869 | 14.056 | 462365 | 286917.4 | 94218.9 | 1.26E+05 | 1.71E+05 | 1.07E+05 | 19460.2 | 34504.5 |
| C19488 | 245.22368 | 18.876 | 23284.2 | 21125.99 | 7442.643 | 10666.95 | 6971.527 | 7803.337 | 66790.8 | 1.07E+05 |
| C19490 | 156.9987 | 1.47 | 62250.2 | 35382.26 | 22016.59 | 25727.08 | 22110.85 | 22948.45 | 203104 | 76757.7 |
| C19529 | 298.24952 | 2.062 | 4.98E+06 | 4.27E+06 | 1.85E+06 | 2.71E+06 | 2.01E+06 | 1.50E+06 | 134010 | 1.64E+05 |
| C19543 | 299.15365 | 16.974 | 23393.3 | 27605.46 | 10774.1 | 15297.26 | 15662.6 | 11131.65 | 220913 | 1.58E+06 |
| C19563 | 258.08641 | 8.772 | 40827.6 | 22082.83 | 2504.637 | 2715.876 | 2596.082 | 2720.138 | 5161.83 | 10741.4 |
| C19566 | 88.99186 | 1.901 | 78333.6 | 31052.37 | 7084.801 | 7756.293 | 7812.208 | 8491.108 | 213533 | 75214.7 |
| C19607 | 172.02116 | 1.687 | 84472.4 | 45765.69 | 11086.67 | 11233.99 | 11036.1 | 10671.94 | 73901.4 | 47989.7 |
| C19615 | 320.0385 | 12.137 | 149336 | 245265.1 | 43218.13 | 3.85E+05 | 4.71E+05 | 3.91E+05 | 118678 | 38090.5 |
| C19620 | 600.24106 | 16.321 | 43999.5 | 73789.34 | 85890.15 | 2.37E+05 | 86869.49 | 1.04E+05 | 64237 | 22842.9 |
| C19670 | 129.04257 | 1.228 | 174382 | 179334.6 | 30201.8 | 28119.83 | 30296.76 | 30263.11 | 671875 | 8.04E+05 |
| C19691 | 335.10753 | 9.678 | 50928.9 | 21460.41 | 2217.144 | 2382.11 | 2298.797 | 2294.232 | 52543.9 | 54146.1 |
| C19757 | 168.11468 | 9.344 | 49751.9 | 40464.64 | 4330.087 | 4868.206 | 4960.323 | 4705.65 | 139504 | 62541 |
| C19891 | 281.61583 | 17.165 | 151676 | 67791.92 | 6463.008 | 7309.564 | 6321.622 | 7159.963 | 44097.9 | 42076.3 |
| C20310 | 330.27633 | 18.754 | 99442.1 | 91236.71 | 18279.97 | 22955.66 | 17929.72 | 22272.89 | 5.10E+06 | 7.02E+06 |
| C20313 | 330.27635 | 19.009 | 99104.6 | 91971.52 | 19078.32 | 23373.7 | 17010.39 | 22491.8 | 528609 | 1.27E+06 |
| C20775 | 421.13103 | 19.452 | 3010.31 | 2675.005 | 2585.114 | 3747.71 | 2658.014 | 4069.602 | 70266.1 | 34657.5 |
| C20792 | 129.04254 | 12.186 | 89599.6 | 55172.56 | 5218.722 | 5874.045 | 5680.957 | 5408.002 | 394170 | 3.54E+05 |
| C20850 | 143.09433 | 1.753 | 118905 | 150025.7 | 1.10E+05 | 1.38E+05 | 1.31E+05 | 1.40E+05 | 1.43E+07 | 1.28E+07 |
| C20988 | 218.06881 | 15.23 | 30738 | 29464.81 | 6140.024 | 8901.861 | 10908.52 | 8537.24 | 100460 | 1.17E+05 |
| C21189 | 210.10078 | 12.276 | 1.09E+07 | 7.74E+06 | 4.12E+06 | 6.23E+06 | 3.49E+06 | 6.42E+06 | 2.10E+06 | 2.87E+06 |
| C22006 | 147.98931 | 1.928 | 3.71E+06 | 2.85E+06 | 1.48E+06 | 1.77E+06 | 2.34E+06 | 1.53E+06 | 2.02E+06 | 1.70E+06 |
| G00093 | 307.22749 | 8.738 | 26837.8 | 23586.7 | 2769.934 | 3001.519 | 2876.598 | 2960.153 | 6091.9 | 12986.9 |
| G00403 | 308.16182 | 16.467 | 30312.2 | 17760.08 | 3317.279 | 3900.566 | 4046.506 | 3177.81 | 11495 | 24023.2 |
| HMDB0001865 | 454.03999 | 14.048 | 1.00E+06 | 1.04E+06 | 5.40E+05 | 9.92E+05 | 1.21E+06 | 7.87E+05 | 23302.1 | 28759.6 |
| HMDB0002513 | 145.05245 | 4.136 | 87989 | 51906.33 | 14785.73 | 16902.99 | 14752.22 | 17442.95 | 1.45E+06 | 9.19E+05 |
| HMDB0002577 | 280.65462 | 20.377 | 51022 | 53034.08 | 11449.32 | 12829.26 | 11492.03 | 13171.93 | 214850 | 3.96E+05 |
| HMDB0004461 | 310.21205 | 11.512 | 12259.2 | 18384.02 | 187822.2 | 181312.6 | 131097.2 | 160973.2 | 32296.2 | 16829 |
| HMDB0005923 | 344.19794 | 1.623 | 259684 | 172189.6 | 3.11E+04 | 3.39E+04 | 3.69E+04 | 3.70E+04 | 195307 | 2.67E+05 |
| HMDB0005960 | 369.16676 | 10.653 | 44108.9 | 64836.8 | 21001.63 | 17695.36 | 20859.21 | 17368.48 | 3.82E+06 | 5.10E+06 |
| HMDB0010221 | 168.11466 | 3.695 | 59821.5 | 52304.87 | 92574.16 | 109147.9 | 42455.09 | 113280.5 | 124654 | 1.04E+05 |
| HMDB0010331 | 144.04187 | 1.058 | 186987 | 145300.2 | 49447.88 | 7.85E+04 | 52970.4 | 4.15E+04 | 2.61E+06 | 3.31E+06 |
| HMDB0010736 | 458.14 | 13.722 | 161989 | 67429.12 | 16667.18 | 26420.66 | 11752.46 | 9114.927 | 73586.9 | 36239.8 |
| HMDB00197 | 682.05293 | 10.193 | 11749.1 | 10085.25 | 47321.05 | 48173.68 | 45713.86 | 48106.05 | 30852.2 | 29023 |
| HMDB00292 | 298.24986 | 15.14 | 176674 | 105158.3 | 8529.322 | 9244.359 | 8681.733 | 8303.52 | 254842 | 1.46E+05 |
| HMDB0062770 | 239.1335 | 10.974 | 69582.2 | 31601.83 | 118432.1 | 100131.3 | 102934.8 | 94200.09 | 252163 | 2.56E+05 |
| HMDB00651 | 155.9047 | 1.855 | 160631 | 103357 | 5.38E+03 | 5.12E+03 | 5.70E+03 | 4748.588 | 338008 | 1.85E+05 |
| HMDB00682 | 328.16533 | 10.763 | 45889.3 | 35323.43 | 2298046 | 2249216 | 1758620 | 2109033 | 43984.9 | 16021.8 |
| HMDB00701 | 803.36661 | 2.313 | 9.87E+06 | 5.62E+06 | 1.59E+05 | 3.52E+05 | 3.62E+05 | 1.25E+05 | 114107 | 1.35E+05 |
| HMDB00705 | 226.15731 | 1.761 | 124744 | 88009.46 | 6.28E+03 | 6.64E+03 | 6.32E+03 | 6.58E+03 | 3.69E+06 | 1.61E+06 |
| HMDB00708 | 197.03779 | 1.437 | 39302.9 | 33425.43 | 91991.35 | 183667.8 | 66692.87 | 131770.3 | 592513 | 5.41E+05 |
| HMDB00832 | 162.08284 | 20.064 | 121582 | 430024.3 | 15326.51 | 1.98E+04 | 27464.67 | 1.41E+04 | 88931.1 | 88975.8 |
| HMDB00888 | 298.25013 | 16.556 | 16900.5 | 28447.08 | 7281.709 | 8159.766 | 7454.404 | 7857.4 | 189403 | 7.85E+05 |
| HMDB01046 | 400.33303 | 16.828 | 159703 | 87314.02 | 236820.4 | 290610.9 | 101184.8 | 573830.6 | 96195.8 | 34832 |
| HMDB01624 |  | 22.16 | 618759 | 1.14E+06 | 8.67E+04 | 1.81E+05 | 2.36E+05 | 1.68E+05 | 136304 | 2.34E+05 |
| HMDB01863 | 540.24513 | 14.516 | 228473 | 485207.6 | 43683.17 | 3.90E+04 | 4.76E+04 | 3.51E+04 | 17329.2 | 12807.7 |
| HMDB02203 | 330.21812 | 10.628 | 141355 | 49911.77 | 8537.566 | 9169.695 | 9290.597 | 9002.066 | 53535.9 | 32843.9 |
| HMDB0244529 | 175.03437 | 12.473 | 83915.1 | 79174.57 | 20577.28 | 22519.78 | 21641.68 | 23212.18 | 1.45E+06 | 1.21E+06 |
| HMDB02833 | 136.06355 | 2.869 | 196338 | 136924.6 | 21555.6 | 21233.9 | 19179.76 | 21130.82 | 93305.5 | 1.11E+05 |
| HMDB04998 | 99.01733 | 1.465 | 110751 | 93412.09 | 104468.6 | 57730.85 | 104058.2 | 63350.29 | 1.44E+07 | 1.14E+07 |
| HMDB05015 | 282.92219 | 22.253 | 71051.7 | 52381.83 | 1.65E+04 | 16323.21 | 1.62E+04 | 16027.92 | 173547 | 1.93E+05 |
| HMDB06029 | 143.09513 | 1.724 | 113757 | 70330.05 | 30145.54 | 33446.47 | 35828.94 | 34642 | 94478 | 84602.6 |
| HMDB10316 | 319.05126 | 17.489 | 155476 | 186980.6 | 858730.2 | 755320.2 | 647391.5 | 778267.1 | 1.11E+07 | 1.34E+07 |
| HMDB13041 | 99.06839 | 2.018 | 1.82E+06 | 1.71E+06 | 8.23E+03 | 7.62E+03 | 8.14E+03 | 7.56E+03 | 5.04E+07 | 5.25E+07 |
| HMDB13677 | 176.10439 | 9.898 | 83170.9 | 24522.52 | 27504.19 | 29413.51 | 27201.74 | 28310.48 | 84111.7 | 44297.9 |
| HMDB13892 | 83.07387 | 1.821 | 300658 | 140884.9 | 8565.894 | 9373.204 | 8839.375 | 9210.161 | 816898 | 8.69E+05 |
| HMDB14375 | 326.14963 | 16.299 | 143141 | 97883.87 | 525158.7 | 564079.4 | 409059.1 | 507587.6 | 111426 | 1.36E+05 |
| HMDB14515 | 164.06795 | 1.732 | 1.12E+06 | 1.75E+06 | 3.02E+04 | 3.11E+04 | 3.35E+04 | 3.18E+04 | 2.01E+06 | 9.93E+05 |
| HMDB14660 | 189.04233 | 12.874 | 200697 | 180125.1 | 111317.3 | 152885.8 | 202345.4 | 139493.5 | 1.27E+08 | 1.37E+08 |
| HMDB14757 | 216.03911 | 1.69 | 345269 | 226193.5 | 1.09E+05 | 2.01E+05 | 1.01E+05 | 1.46E+05 | 3.37E+06 | 1.14E+06 |
| HMDB15316 | 788.43595 | 3.501 | 643327 | 455515.3 | 1.97E+04 | 2.91E+04 | 1.19E+04 | 3.20E+04 | 72965.7 | 63097.6 |
| HMDB15334 | 180.0637 | 5.17 | 30441.6 | 26066.18 | 1983371 | 1908686 | 1960621 | 1747297 | 21593.2 | 18423.6 |
| HMDB15390 | 118.06214 | 3.317 | 4.21E+06 | 3.19E+06 | 1.02E+04 | 1.07E+04 | 1.07E+04 | 1.01E+04 | 527548 | 5.60E+05 |
| HMDB15557 | 363.66826 | 11.998 | 71006.1 | 92128.45 | 9065.956 | 8335.249 | 7825.464 | 7956.266 | 97099.9 | 5.40E+05 |
| HMDB34738 | 258.04002 | 1.946 | 86481.5 | 44171.34 | 17235.56 | 20548.23 | 16566.14 | 20164.85 | 39904.2 | 31884.5 |
| HMDB35724 | 239.21358 | 18.426 | 59536.3 | 85599.12 | 11703.14 | 10913.41 | 12170.12 | 10920.55 | 450853 | 3.10E+05 |
| HMDB36897 | 350.1024 | 21.018 | 113694 | 68884.15 | 6612.32 | 7726.94 | 7006.196 | 7382.051 | 213623 | 14424.7 |
| HMDB38249 | 932.58536 | 16.787 | 137826 | 78372.23 | 243579 | 310774.2 | 310480 | 196725.4 | 128263 | 1.34E+05 |
| HMDB41807 | 404.13197 | 9.687 | 847304 | 512017.4 | 1.79E+05 | 2.97E+05 | 3.09E+05 | 2.70E+05 | 25966.3 | 18682.9 |
| HMDB41889 | 378.09754 | 2.147 | 341592 | 202255.4 | 2.82E+04 | 3.05E+04 | 3.37E+04 | 3.41E+04 | 138597 | 79721.2 |
| HMDB41925 | 226.1676 | 7.526 | 109028 | 86955.24 | 8973.21 | 10405.7 | 9420.654 | 10172.05 | 4.04E+06 | 4.21E+06 |
| HMDB41928 | 226.16759 | 13.868 | 73866 | 50693.07 | 3246.942 | 3636.812 | 4067.762 | 3446.294 | 473371 | 3.16E+05 |
| HMDB59711 | 531.32849 | 14.557 | 36043 | 34002.65 | 12372.27 | 19215.06 | 20826.28 | 15600.8 | 73783.7 | 16125.5 |
| HMDB59876 | 330.27597 | 12.848 | 17439.4 | 25188.64 | 158113.1 | 144749.5 | 152378.9 | 142784 | 5.56E+06 | 5.77E+06 |
| HMDB60906 | 239.08817 | 9.939 | 523481 | 272342.6 | 9.55E+03 | 1.17E+04 | 1.29E+04 | 1.11E+04 | 641238 | 70618.2 |
| HMDB61033 | 716.12868 | 14.193 | 58256.8 | 40252.09 | 9553 | 11716 | 12850 | 11125 | 1.86E+06 | 1.95E+06 |
